# The D2-*mdx* mouse as a preclinical model for Duchenne muscular dystrophy: a natural history study across two independent sites

**DOI:** 10.64898/2026.07.08.737223

**Authors:** Paola Mantuano, Antonietta Mele, Brigida Boccanegra, Christa Tanganyika-de Winter, Davy Van De Vijver, Anne-Fleur Schneider, Marco Mele, Ornella Cappellari, Lisamaura Tulimiero, Sarah Engelbeen, Ernst Suidgeest, Louise van der Weerd, Annemieke Aartsma-Rus, Annamaria De Luca, Heather Gordish-Dressman, Maaike van Putten

## Abstract

**Introduction:** The quality of preclinical studies for rare diseases, such as Duchenne muscular dystrophy (DMD), relies on the availability of comprehensive natural disease history data. In addition to the classic BL10-*mdx* mouse, in recent years, the D2-*mdx* model has increasingly been used as an alternative model due to its reportedly more severely impaired phenotype. To improve our understanding of disease progression in these two DMD models, we conducted a comprehensive natural history study.

**Materials and Methods:** This involved a cross-sectional analysis of key *in vivo* and *ex vivo* outcome measures performed in two independent laboratories, using the same study setup in compliance with TREAT-NMD Standard Operating Procedures (SOPs), while also taking advantage of site-specific expertise. Globally, largely comparable results were obtained across the two study sites.

**Results:** Body composition showed pronounced differences between the strains, with BL10-*mdx* mice displaying a hypertrophic and D2-*mdx* mice displaying an atrophic phenotype. Dystrophic mice of each strain exhibited significant alterations of disease-relevant indices related to muscle functionality and integrity, mostly worsening with age, in comparison to their wildtypes. Cardiac function was affected earlier and more severely in D2-*mdx* mice.

**Discussion:** Notably, for some parameters, genetic-background related differences were observed, emphasizing the need to include control groups with matching genetic backgrounds in experimental designs.

**Conclusions:** Collectively, our natural history study provides benchmark data for these two *mdx* mouse strains to guide model selection for preclinical DMD studies, allowing accurate data interpretation.

**Highlights:**

- **Distinct body composition phenotypes**: BL10-*mdx* mice exhibit pseudohypertrophy while D2-*mdx* mice display pronounced atrophy.
- **Earlier cardiac dysfunction in D2-*mdx***: D2-*mdx* mice develop reduced ejection fraction and stroke volume from 28 weeks, while BL10-*mdx* only at 52 weeks.
- **Genetic background-dependent variations**: Intrinsic deficits in wildtype D2 mice demonstrate that genetic background influences outcome measures independent of dystrophic pathology.
- **Comparable *ex vivo* muscle physiology**: Despite divergent *in vivo* phenotypes, isolated muscle contractile parameters show similar impairment in both dystrophic models.
- **Multi-site standardized validation**: Cross-sectional study at two independent laboratories following harmonized TREAT-NMD Standard Operating Procedures.

## 1. Introduction

Duchenne muscular dystrophy (DMD) is an X-linked, recessive muscle disorder affecting 1 to 5000 live male births (1). *DMD* gene mutations prevent production of functional dystrophin proteins, leading to progressive muscle degeneration and premature mortality in DMD patients (2). Dystrophin is a subsarcolemmal protein associated with the dystrophin-glycoprotein complex (DGC), that ensures a physical linkage between the intracellular cytoskeleton and the extracellular matrix (3). Lack of dystrophin triggers a complex cascade of pathological alterations leading to myofiber degeneration and severe muscle wasting (4,5). Membrane instability, impaired calcium homeostasis, reactive oxygen species overproduction and dysmetabolism, chronic inflammation and fibrosis have been deciphered as crucial events contributing to the progression of pathology in a reinforcing loop ^4,5^. In view of the pathophysiological complexity of DMD, the availability of a reliable animal model capable of providing reproducible and predictive results is crucial, especially when testing potential therapeutics.

The C57BL/10ScSn-*Dmd^mdx^* (BL10-*mdx*) mouse is the genetic and biochemical homolog of human DMD (6). The BL10-*mdx* mouse has been extensively studied and validated over the last 40 years. These mice harbour a spontaneous nonsense point mutation in exon 23 of the *Dmd* gene, leading to a premature stop codon that results in the synthesis of incomplete non-functional dystrophins (7). Loss of dystrophin induces the onset of the typical structural and functional hallmarks of dystrophic pathology (8). Cardiomyopathy appears at a later stage of the disease (around the 8^th^ month of age (9)), whilst extensive fibrosis occurs earlier in the diaphragm, the most affected muscle in BL10-*mdx* mice. Although the BL10-*mdx* mouse allowed gaining insights into pathogenesis and identifying druggable targets, its milder phenotype and the slower disease progression compared to DMD patients (10), question its usefulness in preclinical research. The newly generated D2.B10-*Dmd^mdx^* (D2-*mdx*) mouse, obtained by crossing classic BL10-*mdx* with DBA/2J wildtype (D2-WT) mice, seems to exhibit a more severe dystrophic phenotype that is claimed to better recapitulate human pathology and to improve data translatability (11,12). Indeed, this model has a polymorphism in the *Ltbp4* gene, that increases TGF-beta signalling, similar to the detrimental *LTBP4* allele seen in DMD patients (13,14). Consequently, an exacerbation of inflammatory and pro-fibrotic events has been described in D2/*mdx* mice (11,15). Furthermore, the presence of the *Dyscalc* locus of either the *Abcc6* or *Emp3* gene is thought to be responsible for calcifications observed in skeletal and cardiac muscle of D2-*mdx* mice (11,16), while a splice site variant in the *Anxa6* gene likely underlies an impairment of the self-renewal activity of satellite cells, worsening their muscle-repairing capacity (12,17). Earlier signs of functional and morphological cardiac compromission have been found in D2-*mdx* mice (11).

In spite of this seemingly more severe phenotype, a complete picture of the natural history of the D2-*mdx* phenotype is missing, while fluctuations of the disease signs over life span have been described (18). In parallel, the use of the D2-*mdx* mouse for drug testing is progressively increasing (19). In this framework, the project entitled “Of Mice and Measures” (OMAM) aimed to bridge this gap (20). This effort consists of a cross-sectional study performed at two independent study sites that followed TREAT-NMD Standard Operating Procedures (SOPs) to gain a better understanding of the D2-*mdx* model in comparison with the classic BL10-*mdx* mouse. The main goal is to assess, by means of an independent validation, the predictivity of the D2-*mdx* model, the most appropriate time window to conduct preclinical studies in relation to the disease progression and the best model for proof-of-concept studies based on drug mechanism of action. With this we aim to contribute to the advancement of translatability of preclinical findings to the clinic. In this study, we performed a comprehensive set of key *in vivo,* and *ex vivo* analyses with outcome measures assessing skeletal and cardiac muscle functionality. These were independently performed by two institutes by means of identical or comparable techniques, additionally taking advantage of site-specific expertise. Our results provide the DMD field with benchmark data for the D2-*mdx* and BL10-*mdx* mouse models.

## 2. Materials and Methods

### 2.1 Animal care

At the University of Bari “Aldo Moro” (UniBa), 42 male mice *per* genotype (C57BL/10ScSn/J wildtypes (BL10-WT), C57BL/10ScSn-*Dmd^mdx^*/J (BL10-*mdx*), DBA/2J (D2-WT) and D2.B10-*Dmd^mdx^*/J (D2-*mdx*)) of 4 weeks of age were purchased from The Jackson Laboratory (USA, distributed by Charles River, Calco, Italy) to perform the study. All mice were housed in suitable cages (max. 5 mice *per* cage) filled with sawdust and bedding (LIGNOCEL 3-4S, Charles River Laboratories) and acclimatized to local housing conditions (22 – 24°C, humidity 50 – 60%, and 12 h light/12 h dark cycle) for approximately 5 days. All mice were got access to a daily amount of chow (RM3, Special Diets Services (SDS), Essex, United Kingdom) of 5 g *per* mouse. This fixed amount, largely exceeding the physiological daily consumption (∼3 g/mouse) allowed proper control by the experimenter of any possible macroscopic deviation in food consumption between groups, which in fact did not occur. Filtered tap water was available *ad libitum*. All experiments were conducted in conformity with the Italian Guidelines for Care and Use of Laboratory Animals (D.L.116/92), and with the European Directive (2010/63/UE). The study has been approved by the National Ethics Committee for Animal Research & Welfare of the Italian Ministry of Health (authorization N. 1119/2020-PR).

At the Leiden University Medical Center (LUMC), breeding couples of the four strains were obtained from The Jackson Laboratory, and their offspring, generated at the in-house breeding facility of the LUMC, was used for the experiments. Fifty males *per* genotype (BL10-WT, BL10-*mdx,* D2-WT and D2-*mdx*) were generated for experiments. Mice were housed in groups of 2 to 5 animals in individually ventilated cages (Makrolon type II) filled with sawdust and enriched with nesting (Bed-r’Nest BRN8SR) and bedding (LIGNOCEL BK-8-15-00433) materials and a cardboard tunnel (GLP fun tunnels mini 1022006). *Ad libitum* access to water and standard RM3 chow (SDS, Essex, United Kingdom) was provided to the mice. All tests were performed in rooms dedicated to behavioural experiments and timing of each test was kept as consistent as possible for all the animals. The experiments were approved by the Animal Ethics Committee of the LUMC (AVD 1160020171407, PE.17.246.037) and executed conform with the Directive 2010/63/EU of the European Parliament.

### 2.2 Experimental groups

At each study site, mice were randomized over several experimental cohorts, which underwent identical parallel assessments, differing primarily in the time-point chosen for sacrifice. Mice were either sacrificed at 2 (LUMC only), 4, 8, 12, 28, or 52 weeks of age (Figure 1). Cohorts consisted of *n* = 8 males *per* mouse strain, *per* study site, except for those sacrificed at 52 weeks of age, which contained two additional mice to accommodate for potential unforeseen deaths. Body weight (BW) and forelimb grip strength was assessed monthly by both sites, except for cohorts sacrificed at 2 or 4 weeks of age for which only an end-point measurement was performed. At LUMC, body composition, wire hanging performance, spontaneous activity and respiratory functionality were assessed on a monthly basis. At UniBa, hindlimb plantar flexor torque and ultrasonography of the hindlimb and diaphragm was performed in mice aged 8, 12, 28, and 52 weeks; specifically, torque was measured in all mice regardless of the cohort at these ages until sacrifice at the respective endpoint, while ultrasound measurements were performed as a terminal assessment. Furthermore, functionality of the heart was assessed for mice aged 8, 12, 28, and 52 weeks (ultrasonography; UniBa, magnetic resonance imaging; LUMC), while markers for muscle integrity were assessed for all cohorts. Lastly, *ex vivo* physiology of the extensor digitorum longus (EDL) and diaphragm was assessed at UniBa after sacrifice, at the same endpoints.

**Figure 1.**
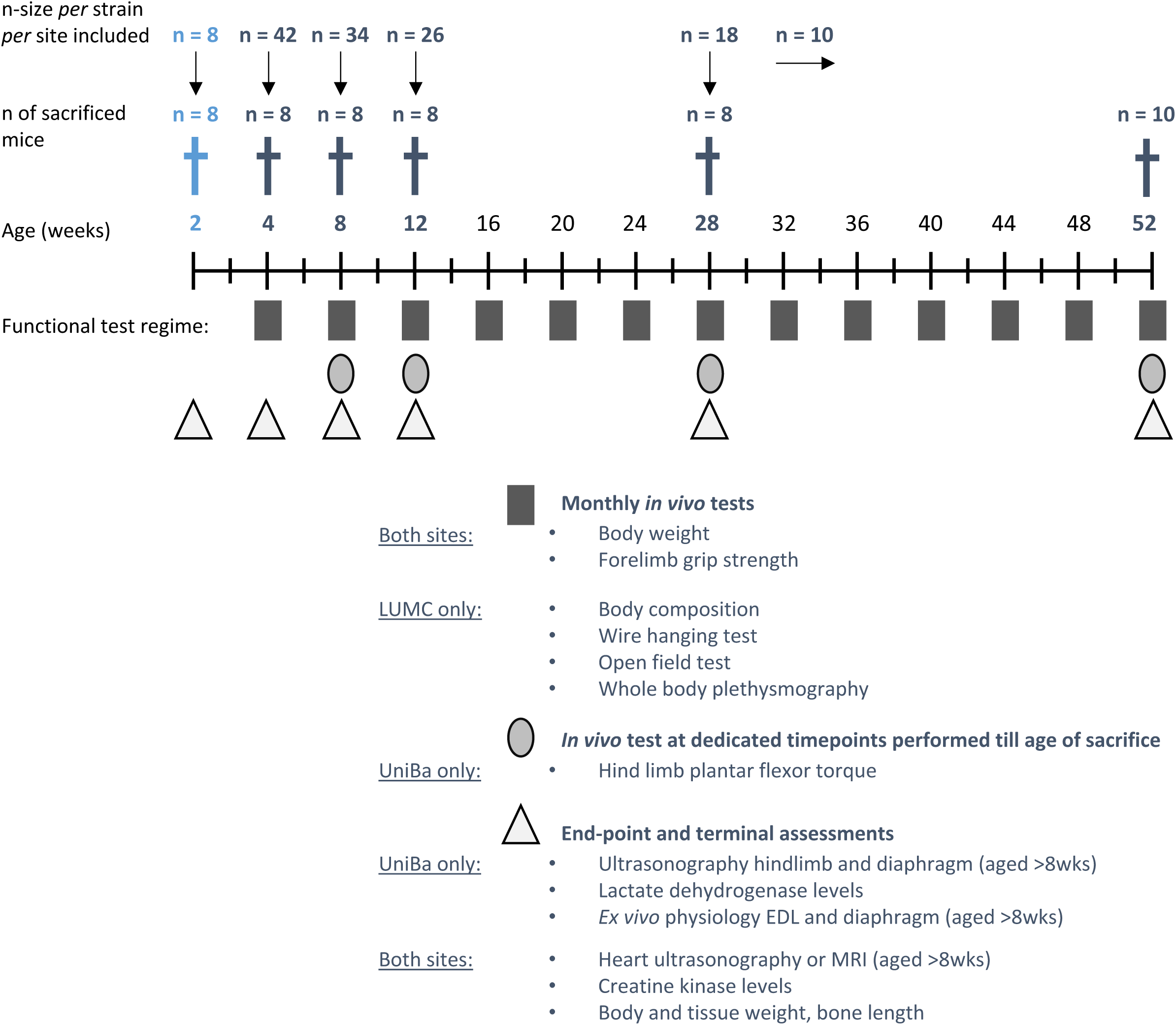
Study design. Animals were subjected to a cross-sectional study at two independent study sites. Assessments were either performed by both sites (body and muscle weight, forelimb grip strength, cardiac function, creatine kinase levels, and bone length), or by a single site (LUMC; body composition, wire hanging test, open field test, and whole-body plethysmography. UniBa; Hindlimb plantar flexor torque, ultrasonography of the hindlimb and diaphragm, lactate dehydrogenase levels, and *ex vivo* physiology of the EDL and diaphragm). Mice belonging to the 2 and 4-week cohorts only underwent tissue weights and plasma biomarker assessments. Torque measurements were performed in mice from 8 weeks onwards. Two-week-old mice (light blue) were only included at LUMC. N; number, EDL; extensor digitorum longus, MRI: magnetic resonance imaging. LUMC: Leiden University Medical Center, UniBa; University of Bari.

Mice were sacrificed by cervical dislocation (or decapitation for those aged 2 weeks). Finally, forelimb and hindlimb muscles (quadriceps; gastrocnemius; tibialis anterior; triceps, and EDL of both limbs), the heart and vital organs (liver, kidneys, and spleen) of all mice were weighed, and the average weight was recorded. Lastly, length of the tibia and/or femur bones was measured.

### 2.3 Body composition

To assess body composition of the mice, lean and fat mass was measured with the EchoMRI-100 body composition analyzer (EchoMRI, Houston, Texas, USA) on a monthly basis at LUMC. Prior to the measurements, a system test was performed using canola oil. Mice were weighed, placed inside the tube and inserted in the EchoMRI. Both absolute values of lean or fat mass divided by BW were used for analyses.

### 2.4 Forelimb grip strength

At both study sites, forelimb grip strength was measured on a monthly basis by means of an isometric force transducer (Columbus Instruments, USA) according to a standard protocol (TREAT-NMD SOP (ID) Number: DMD_M.2.2.001). In detail, the mouse was suspended by the tail and moved horizontally towards a metal triangle allowing it to grasp it with its forelimbs. The operator pulled the mouse away until its grasp was broken: the force produced was measured by the force transducer attached to the triangle. The test was repeated 5 times with a 1-minute break between measurements. For data analysis, we used the maximal value obtained *per* mouse (absolute maximal force, kg force, kgF) and normalized to BW (kgF/kg).

### 2.5 Wire hanging test

At LUMC, hanging performance was assessed with the wire hanging test according to previously described protocols and TREAT-NMD SOP DMD_M.2.1.004, once monthly. The mouse was handled by its tail and brought near the metal wire that was placed 37 cm above a large cage filled with bedding. When it grasped the wire with its two forepaws, it was released and the timer was started. The use of the hindlimbs and tail was allowed during hanging and primarily observed in strong wildtype animals. The timer was stopped when the mouse fell off the wire or when the maximum hanging time of 10 minutes was reached. Animals that failed to hang for 10 minutes were given a maximum of two additional attempts to do so. The longest hang time achieved was used for analyses.

### 2.6 Open field test

At LUMC, spontaneous activity was assessed with the open field test once a month. Mice were released in the middle of a white squared box (50 x 50 x 35 cm) and allowed to freely move around for 10 minutes, while being video recorded by an overhead camera. The box was cleaned with 70% ethanol after each trial to remove odor markings. For analysis, the box was digitally divided into an inner (region of 30 x 30 cm in the middle of the box) and an outer zone (the remaining area). Movement and location were automatically measured after tracking with the Ethovision XT Software at a rate of 20 frames per second. Quality of the tracking of the mice was confirmed by checking the duration of untracked video. Per trial, on average 3.62 seconds were not tracked, except for four videos in which tracking was unsuccessful for 33 to 53 sec. We excluded four trials in which >1 minute of video could not be correctly tracked due to overexposure (three D2-*mdx* mice aged 4 weeks and one D2-*mdx* mouse aged 44 weeks).

### 2.7 Respiratory function analysis

At LUMC, respiratory function of the mice was assessed using the whole-body plethysmography (RM-80; Columbus Instruments) on a monthly basis. Hereto, mice were weighed and left to acclimatize in the setup for a minimum of 30 seconds before respiration was recorded using the Axoscope software version 10.4 (Molecular Devices, San Jose, CA, USA) for two minutes. Respiration rate and amplitude were analyzed with the event detection feature of the Clampfit 10 program (Molecular Devices).

### 2.8 Hindlimb plantar flexor torque

The isometric torque produced by hindlimb plantar flexors (gastrocnemius, soleus – SOL, and plantaris muscles) was longitudinally measured by means of the 1300A 3-in-1 Whole Animal System (Aurora Scientific Inc. – ASI, Aurora, ON, Canada). Mice underwent inhalation anesthesia (∼3% isoflurane in an induction chamber, then ∼2% isoflurane via nose cone for maintenance, both with 1.5 L/min O_2_), delivered via a vaporizer (Harvard Apparatus Fluovac and Datex Ohmeda Isotec 4, Holliston, MA, USA) with an oxygen concentrator (LFY-1-5A, Longfei Group Co., Wenzhou, China; distributed by 2Biological Instruments, Besozzo, VA, Italy).

After prepping the skin of the right hindlimb by removing hair and cleaning, the animal was placed supine on a temperature-controlled platform (mod. 809B, ASI) set at 36°C; the right paw was taped to a footplate connected to a dual-mode servomotor (mod. 300C-LR, ASI), forming a 90° angle with the hindlimb secured at the knee. Contractions were elicited via percutaneous electrical stimulation of the sciatic nerve, through a pair of needle electrodes (Chalgren Enterprises Inc., Gilroy, CA, USA) connected to a high-power, bi-phase stimulator (mod. 701C, ASI), in turn controlled by a data acquisition signal interface (mod. 604A, ASI) and by ASI Dynamic Muscle Control software (DMCv5.415). Initial twitches, elicited using 0.2 ms single square wave pulses, were used to adjust the current (ranging from 25 to 40 mA) to optimize torque production. After this, a series of isometric contractions were evoked at increasing frequencies (200 ms pulses at 1, 10, 30, 50, 80, 100, 120, 150, 180, and 200 Hz, one every 30 s). Data for plantar flexor torque (N*cm) recorded at each frequency were obtained via ASI Dynamic Muscle Analysis software (DMAv5.201), normalized to each mouse BW (N*mm/kg), and used to construct torque – frequency curves (21–24).

### 2.9 Ultrasonography of hindlimb, diaphragm, and heart

At UniBa, ultrasonography evaluations were performed on hindlimb, diaphragm, and heart for the mice cohorts of 8, 12, 28 and 52 weeks prior to sacrifice. To avoid additional stress to the animals that could influence the outcome of the procedure, an interval of more than 48 hours was maintained between the final *in vivo* test and the ultrasound assessments. Ultrasonography experiments were carried out by means of the ultra-high frequency ultrasound biomicroscopy VisualSonics Vevo® 2100 Imaging System (VisualSonics, Toronto, ON, Canada), which allows multiple image acquisition modes (25). According to TREAT-NMD SOP DMD_M.2.2.003, the animals were properly prepared prior to the imaging session to allow optimal image acquisitions. Each mouse was anesthetized via inhalation of isoflurane (induction with ∼3% isoflurane and 1.5% O_2_ l/min, then constantly maintained at ∼2% isoflurane and 1.5% O_2_ l/min via a nose cone) and placed on a thermostatically controlled table (kept at 37°C). This latter was equipped with 4 copper leads which allowed the monitoring of both heart and respiratory rate of the mouse to minimize any physiological variation. Body temperature was monitored during the imaging session using a rectal probe. A petroleum-based lubricant was used to protect the eyes from drying. Fur from the thorax and hindlimbs was removed using depilatory cream to avoid interference during image acquisition, and preheated ultrasound gel was applied between the skin and the probe to ensure proper image recording.

#### 2.9.1 Diaphragm ultrasonography

As described previously (26,27), each mouse was placed in a dorsal decubitus position, and the platform was tilted by 30° from horizontal with the head of the mouse lower than the feet. The probe was positioned transversally to the midsternal of the mouse at 120° with respect to the platform. Images were acquired in one-dimensional (M-Mode) and two-dimensional (B-Mode), using a probe operating at a frequency of 21 MHz characterized by a lateral and axial resolution of 165 and 75 μm, respectively. Movement amplitude was measured in M-mode during normal breathing cycles on the left side, which provides less variability in measurements for both WT and dystrophic mice. The amplitude during each inspiration (positive deflection) was measured as the distance (in mm) between the baseline and the peak of contraction. For each mouse, the amplitude was calculated as the mean value obtained from 3-5 measurements. The images acquired in B-mode were used to assess the echodensity of the diaphragm, which was measured using ImageJ® software by creating a greyscale analysis histogram over the entire section of the diaphragm delineated with a constant size of 4514.0 ± 17.6 pixels. For each mouse, echodensity was obtained as the main value from 4 frames of the same acquisition drawing the region of interest (ROI) in the same area of the diaphragm. For each strain, variations in echodensity were expressed as percentage differences between WT and *mdx* groups of the mean echo intensity of the pixels included in the outlined area.

#### 2.9.2 Hindlimb ultrasonography

For these measurements (25), each mouse was placed in a ventral decubitus position. The hindlimbs were positioned closely parallel to the body, with each foot forming a 90° angle with the hindlimb. After applying the ultrasound gel, a three-dimensional (3D) volumetric scan of the hindlimbs was acquired by translating the ultrasound probe parallel to the long axis of the hindlimb. Multiple two-dimensional (2D) images were then acquired at regular intervals in Power Doppler mode by using an MS250 probe at a frequency of 21 MHz. At the end of the procedure, 3D images were reconstructed from multiple 2D frames previously collected and visualized using VisualSonics 3D software. This allowed to calculate the total hindlimb volume (in mm^3^). The 2D images were used to evaluate the echodensity of the gastrocnemius muscle – selected as a representative example of a muscle severely affected by dystrophic pathology and easily identifiable in ultrasonographic images – via ImageJ® similarly to what was described for the diaphragm. Specifically, the whole muscle was outlined by obtaining a ROI section of constant size of 3187.8 ± 118.2 pixels. Again, for each strain, changes in echodensity were expressed as percentage differences between WT and *mdx* groups of the mean echo intensity of the pixels included in the delineated area (22–24,28).

#### 2.9.3 Heart ultrasonography

Echocardiography was performed in B-Mode and M-Mode, by using a high-resolution transducer at a frequency of 40 MHz (MS550). Images were acquired in a modified parasternal long axis (PLAX) view, with the animal placed in supine position. Briefly, the left ventricle (LV) M-Mode trace was used to measure stroke volume (SV, µl; blood volume pumped from the LV per beat), cardiac output (CO, ml/min, blood volume pumped per minute), ejection fraction (EF, %; volumetric percentage of blood pumped from LV per beat) and shortening fraction (SF, %; size reduction of ventricular diameter during systole) (29,30).

At the end of the procedure, the anaesthetic influx was discontinued, and each animal was first cleaned with water, returned to its cage, warmed up with a heat lamp to restore the correct body temperature, and monitored until complete recovery.

### 2.10 Cardiac magnetic resonance imaging

At LUMC, cardiac magnetic resonance imaging (MRI) experiments were performed using a 7T PharmaScan with the TX/TR Cryoprobe, with the ParaVision 360 V2.0 software (Bruker BioSpin, Germany), in prone position. Mice were anesthetized by inhalation of 2% isoflurane in a 1:1 mixture of pure oxygen and air. The respiration rate was monitored using a respiration pad (SA Instruments, USA). The body temperature was recorded with a rectal probe and controlled using a thermocouple pad (Medres GMBH, Germany). After acquisition of a localizer scan, cardiac function was assessed on short-axis images which were acquired with the retrospectively-gated Intragate CineFLASH sequence with the following settings: echo time; 1.7ms, repetition time; 67 ms, image matrix size; 200×160, field-of-view; 25×20mm. Ten slices were used to cover the heart and the navigator package was placed parallel to the slices. In-house developed Mass Research Software (31) was used for quantitative assessment of LV function. The result of automated AI-based segmentation of the LV endocardial and epicardial contours was visually reviewed and manual corrections were applied if deemed necessary. End-systolic and end-diastolic cardiac phases were automatically determined and, as quality criterion of the cardiac contouring, a maximum mass difference of the LV of 10% was allowed between both cardiac phases. End-systolic volume (ESV) and end-diastolic volume (EDV) were determined, and heart rate (HR) was extracted from the IntraGate sequence. These parameters were used to calculate the stroke volume (SV□=□EDV□−□ESV), cardiac output (CO□=□SV□×□HR) and ejection fraction (EF□=□SV/EDV□×□100%).

### 2.11 Ex vivo physiology of the diaphragm and EDL

Mouse cohorts underwent *ex vivo* functional tests on isolated diaphragm and EDL muscles at 8, 12, 28, or 52 weeks of age. Mice were anesthetized via intraperitoneal injection with ketamine (100 mg/kg) and xylazine (16 mg/kg), with an additional ketamine boost (30 mg/kg) if needed.

A strip of the right hemidiaphragm (no more than 4 mm wide) was cut from the excised muscle and then tied tightly to the rib and central tendon using a 4/0 suture thread, whilst the EDL muscle of the left hindlimb was tied firmly with 6/0 silk suture thread (Fine Science Tools Inc., Foster City, CA, USA) at the proximal and distal tendons during dissection. Two loops with sutures at both ends were made to place each muscle in a recording chamber containing 25 mL of isotonic Ringer’s solution (21), bubbled with a mix of Clioxicarb (95% O_2_ plus 5% CO_2_), at pH 7.2–7.4 and kept at 27 ± 1°C via a circulating thermostat (Julabo GmbH, Seelbach, Germany). Then, the diaphragm strip was placed into a vertical muscle bath (mod. 800A, ASI), with the central tendon fixed to a hook at the bottom of the chamber, and the rib fixed to a dual-mode muscle lever (mod. 300C-LR, ASI). Similarly, the EDL muscle was placed into a horizontal muscle bath (mod. 809B-25, ASI), with the proximal tendon fixed to a 300C-LR force transducer and the distal tendon fixed to a hook at the opposite side of the chamber. In each bath, electrical field stimulation was obtained by two axial platinum electrodes closely flanking the muscle, connected to a high□power bi□phase stimulator (mod. 701C, ASI). Each apparatus was equipped with a data acquisition signal interface (mod. 604A, ASI) and software (for diaphragm: DMCv4.1.6; for EDL: DMCv5.415, ASI).

After equilibration (∼30 min), each muscle preparation was stretched to its optimal length (L_0_, measured with an external caliper). Single twitch (Ptw) force and contraction kinetics (*i.e.*, time to peak, TTP; half-relaxation time, HRT) were obtained as mean values from 5 twitches elicited by pulses of 0.2 ms, every 30 s. Tetanic contractions were elicited by applying trains of 2.0 ms pulses for 450 ms (diaphragm) or 350 ms (EDL) at increasing frequencies (from 10 to 250 Hz), every 2 min. Maximal tetanic force (P0) was usually recorded at 140–180 Hz.

Following the isometric stimulation protocol, both muscles were subjected to a series of 10 eccentric contractions, every 30 s. Briefly, an initial 300 ms isometric contraction was elicited, followed by a stretch of 10% L_0_ at a speed of 1 L_0_ s^−1^ imposed for the last 200 ms. The progressive decay in isometric force at 5^th^ and 10^th^ pulses was calculated as the percentage of reduction in force vs. the 1^st^ pulse. Two tetanic stimuli (120 Hz, 500 ms) were elicited 5 and 15 min after the eccentric protocol, to calculate the recovery from the stretch-induced force drop vs. the tetanic force registered before the protocol started, as well as muscles’ compliance to stretch.

Results were analyzed via ASI software (DMAv3.2 for diaphragm and DMAv5.201 for EDL), to calculate Ptw, TTP, HRT, and P0. Absolute Ptw and P0 were normalized to muscle cross sectional area according to the equation sP = P/(Mass/L_f_*D) where P is the absolute tension, Mass is the muscle mass, D is the density of skeletal muscle (1.06 g/cm^3^), L_f_ was obtained by multiplying L_0_ by previously determined muscle length to fiber length ratio (diaphragm = 1; EDL = 0.44) (21,22,32).

### 2.12 Plasma creatine kinase and lactate dehydrogenase assessment

At UniBa, blood samples were collected at sacrifice via cardiac puncture of the left ventricle with a heparinized insulin syringe and collected in heparinized tubes. The samples were processed within 30 minutes after collection. Platelet-poor plasma was obtained after a first centrifugation step (20 mins at 4000 rpm at 4°C) to remove almost all blood cells; a second centrifugation step (10 minutes at 12000 rpm) was performed to ensure complete platelet removal. The fresh plasma was used to quantify creatine kinase (CK) and lactate dehydrogenase (LDH) enzymatic activity (U/L) with commercially available diagnostic kits (CK NAC LR and LDH LR, SGM, Rome, Italy). The assays were carried out with a spectrophotometer (Ultrospec 2100 Pro UV/Visible, Amersham Biosciences, Little Chalfont, UK) set to a wavelength of 340 nm at 37°C, according to the manufacturer’s instructions (22).

At LUMC, blood samples were collected, via an angled tail-vein cut, in heparin-coated capillary collection tubes (microvettes CB300 (Sarstedt, Nümbrecht, Germany)), prior to sacrifice. Tubes were stored on ice for a maximum of 2 hours, after which they were centrifuged at 4°C at 13000 rpm for 5 minutes to obtain plasma. CK levels were determined using CK test strips in the Reflotron plus system (Roche, Basel, Switzerland).

### 2.13 Statistical analysis

All statistical analyses were performed using STATA V18 (College Station, TX). A *P*-value of ≤0.05 was considered statistically significant.

#### 2.13.1 Assessment of normality

Outcomes were assessed for normality within each genotype and age group using both a formal Shapiro-Wilk normality test and a visual inspection of histograms. All outcomes, except CK and LDH levels, were shown to be normally distributed, therefore parametric tests were used throughout the analysis. For CK and LDH levels, which did show a departure from normality, individual age comparisons were performed using nonparametric tests and the fit of the longitudinal regression models was verified through regression diagnostics.

#### 2.13.2 Presentation and analysis of observed data

In order to describe the data collected as part of this study, observed data points for each outcome are presented for each of the two study sites independently. Data is presented graphically, either as a line graph (with the 95% confidence intervals as error bars) or as a box plot highlighting each of the four genotypes at each time point assessed. Targeted comparisons were performed among genotypes at each time point where the following genotypes were compared: BL10-WT versus BL10-*mdx*, D2-WT versus D2-*mdx*, BL10-*mdx* versus D2-*mdx* and BL10-WT versus D2-WT. Comparisons of observed values were performed using either a student’s t-test or a Wilcoxon rank sum test and the resulting *P*-values adjusted appropriately for multiple comparisons within each age group using the Sidak method. Those comparisons with an adjusted *P*-value less than 0.05 are indicated symbolically on the figures. In addition, the exact adjusted *P*-values are shown in the Supplementary Tables. 1.

#### 2.13.4 Presentation and analysis of predicted data

In order to show what could be expected from an external experiment in which outcome measures are evaluated using the same validated procedures and experimental conditions, we have used the data observed in this study to generate predicted values of each outcome for each genotype over a range of age groups. We combined observed data points from both sites and used a linear regression model for outcomes assessed only once per mouse (*e.g.,* cardiac functionality, tissue weights) or a mixed effects linear regression model for outcomes assessed multiple times per mouse (BW and forelimb grip strength). We used these models to generate predicted means and 95% confidence intervals at ages 2 to 52 weeks. If the experiment was repeated on additional mice from the same population, the 95% confidence interval indicates the range where we expect the mean to lie. Each model included the outcome as the dependent variable and genotype and age as the independent variables. For the mixed effects models, an additional random term was included to account for the repeated measures. In each model, an interaction between genotype and age was assessed and retained in the model if significant.

## 3. Results

### 3.1 BL10-mdx mice are hypertrophic while D2-mdx mice are atrophic

To gain insight into the natural disease history of the D2-*mdx* mouse, in a direct comparison with the classic BL10-*mdx* strain, a cross-sectional study was performed at the two independent study sites. Muscle functionality and integrity was assessed on a monthly basis, and mice were sacrificed at either 2, 4, 8, 12, 28 or 52 weeks of age (Figure 1). No signs of discomfort (lack of appetite, body weight, or hair loss, stereotypic or aggressive behaviour, etc.) or macroscopic alterations of vital functions were observed in the vast majority of mice throughout the study. At UniBa, nine out of 168 mice died prematurely during the study (12 weeks cohort: 2 BL10-*mdx*, 28 weeks cohort: 1 BL10-*mdx,* 52 weeks cohort: 2 BL10-*mdx,* 3 D2-*mdx* and 1 D2-WT mouse), while at LUMC, five out of 200 mice died prematurely (28 weeks cohort: 1 BL10-WT, 52 weeks cohort: 1 BL10-*mdx*, 1 D2-*mdx*, and 2 D2-WT mice).

Body weight (BW) was assessed on a monthly basis (Figure 2A and Supp Tables 1, 2). Regardless of the study site and strain, all mice showed a physiological and progressive gain in weight over time. In detail, BL10-*mdx* mice were overall heavier than BL10-WT mice up to the age of 28 weeks. Contrastingly, D2-*mdx* mice were lighter than D2-WT and BL10-*mdx* mice throughout the study. These observations are in line with the previously reported pseudohypertrophy (BL10-*mdx*) and atrophy (D2-*mdx*) (33–35). At UniBa, D2-WT mice were consistently lighter than BL10-WT mice, while at LUMC this was clearly observed only up to 16 weeks of age. Overall, mice were heavier at UniBa than at LUMC, resulting in a significant difference between the sites (*P* = 0.034, Supp Table 1). This could have resulted from the more extensive *in vivo* test regime at LUMC, or differences in housing conditions. The greater BW of BL10-*mdx* mice resulted from their substantial lean mass which exceeded that of the other three groups (Figure 2B-C, Supp Table 3). BW-normalized lean mass decreased with age in both BL10 strains, being most pronounced in the wildtypes, whereas it remained stable in adult D2-*mdx* and D2-WT mice. D2-*mdx* mice had a lower absolute lean mass compared to D2-WT mice, but not when corrected for BW. In contrast to the lean mass, BL10-*mdx* mice had limited amounts of fat compared to the other strains throughout the study, being 2.6 times lower than BL10-WT at 52 weeks of age (Figure 2D-E, Supp Table 3). The fat mass of dystrophic and wildtype D2 mice was more comparable but also had a larger interindividual variability. For all parameters (Figure 2A-E, right panel), we observed predicted values that would be expected from an independent experiment conducted under identical experimental conditions.

**Figure 2.**
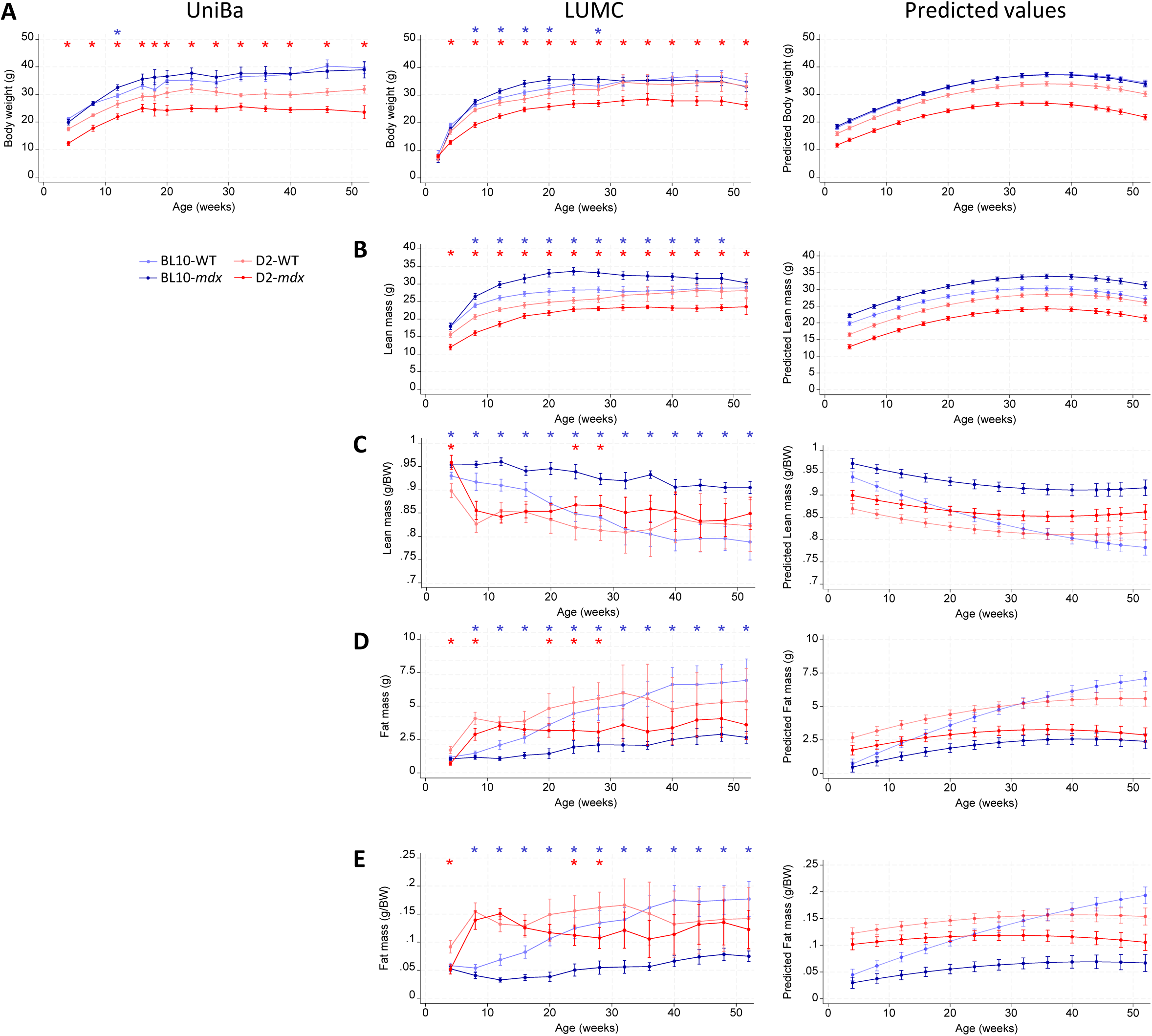
Body weight and composition. Absolute and predicted values are provided for the body weight (A), lean mass (B), body weight normalized lean mass (C), fat mass (D) and body weight normalized fat mass (E). Blue and red stars indicate a significant difference (*P*<0.05) between BL10-WT and BL10-*mdx* or D2-WT and D2-*mdx* mice respectively. UniBa; University of Bari, LUMC: Leiden University Medical Center.

The average weight of several muscles, heart, and organs was determined after sacrifice. Data for absolute and BW-normalized weights and their predicted values are shown in Figure 3, Supp Figures 1, 2 and Supp Tables 4, 5. Both study sites consistently observed that BL10-*mdx* muscles were markedly hypertrophic, in particular from the age of 8 weeks onwards, whilst D2-*mdx* muscles exhibited a progressive atrophy (Figure 3A-D). Normalized heart weights of D2 mice exceeded those of BL10 mice throughout the study at UniBa, and only in mice at later time points (28 and 52 weeks of age) at LUMC. Hearts of both DMD models were heavier compared to their wildtypes at 52 weeks of age at UniBa (Figure 3E). Alterations in organ weights were modest. Kidneys of D2 mice were larger than those of BL10 ones. While the weight did not differ between BL10-*mdx* and BL10-WT mice, kidneys of D2-*mdx* mice were heavier than those of D2-WT mice when normalized to BW (Supp Figure 2A). In parallel, normalized liver weight was generally higher in both DMD strains than in the wildtypes (Supp Figure 2B). Normalized spleen weights and bone length (femur, tibia) were largely comparable between the strains (Supp Figure 2C-E). Study site comparisons of muscle and organ weights revealed differences of small magnitudes for some of the data, reaching significance due to the large sample size (354 samples, Supp Table 1).

**Figure 3.**
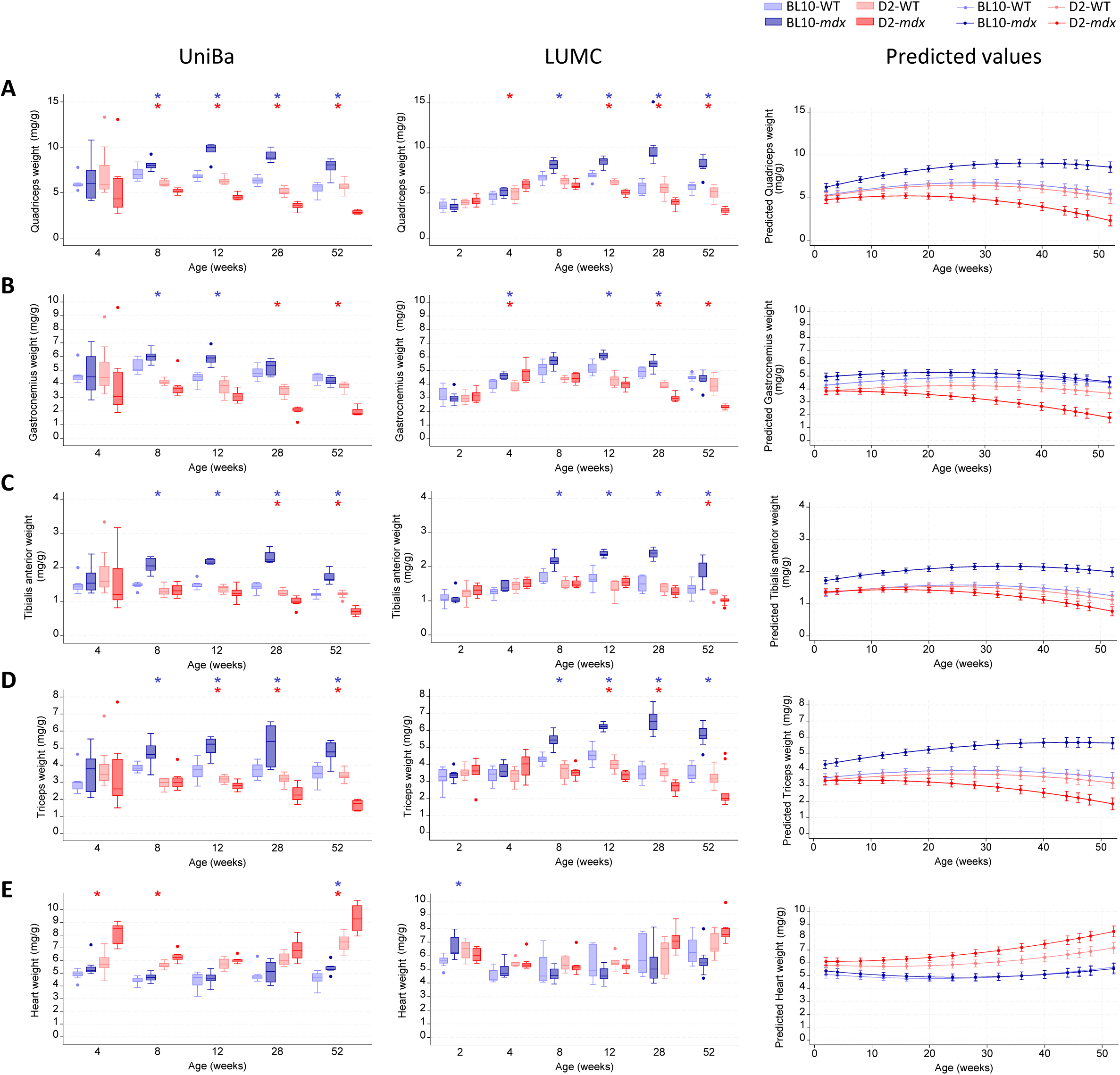
Muscle weights normalized by body weight. Weight analyses corrected for body weight shown as absolute and predicted values for the quadriceps (A), gastrocnemius (B), tibialis anterior (C), triceps (D) and heart (E). Blue and red stars indicate a significant difference (*P*<0.05) between BL10-WT and BL10-*mdx* or D2-WT and D2-*mdx* mice respectively. UniBa; University of Bari, LUMC: Leiden University Medical Center.

### 3.2 Both DMD models display altered in vivo indices associated with muscle function and integrity

Various *in vivo* outcome measures, *i.e.*, grip strength test (UniBa, LUMC), hanging wire, open field, and whole-body plethysmography (LUMC), as direct or indirect indices of muscle function and integrity, were assessed in mice from the age of 4 weeks, until sacrifice.

Both DMD models exhibited reduced absolute and BW-normalized forelimb grip strength values compared to their corresponding wildtype controls (Figure 4A-B and Supp Table 6). D2-*mdx* mice were weaker than BL10-*mdx* mice, when compared in terms of absolute force, only at UniBa. At UniBa, normalized force values of BL10-*mdx* and D2-*mdx* mice nearly overlapped, while at LUMC D2-*mdx* values regularly exceeded those of BL10-*mdx* mice, likely in relation to the differences in BW values observed across the two sites. No differences were, however, found between the two sites in the absolute or BW-normalized maximal force values (*P* = 0.8 and *P =* 0.2 respectively; Supp Table 1).

**Figure 4.**
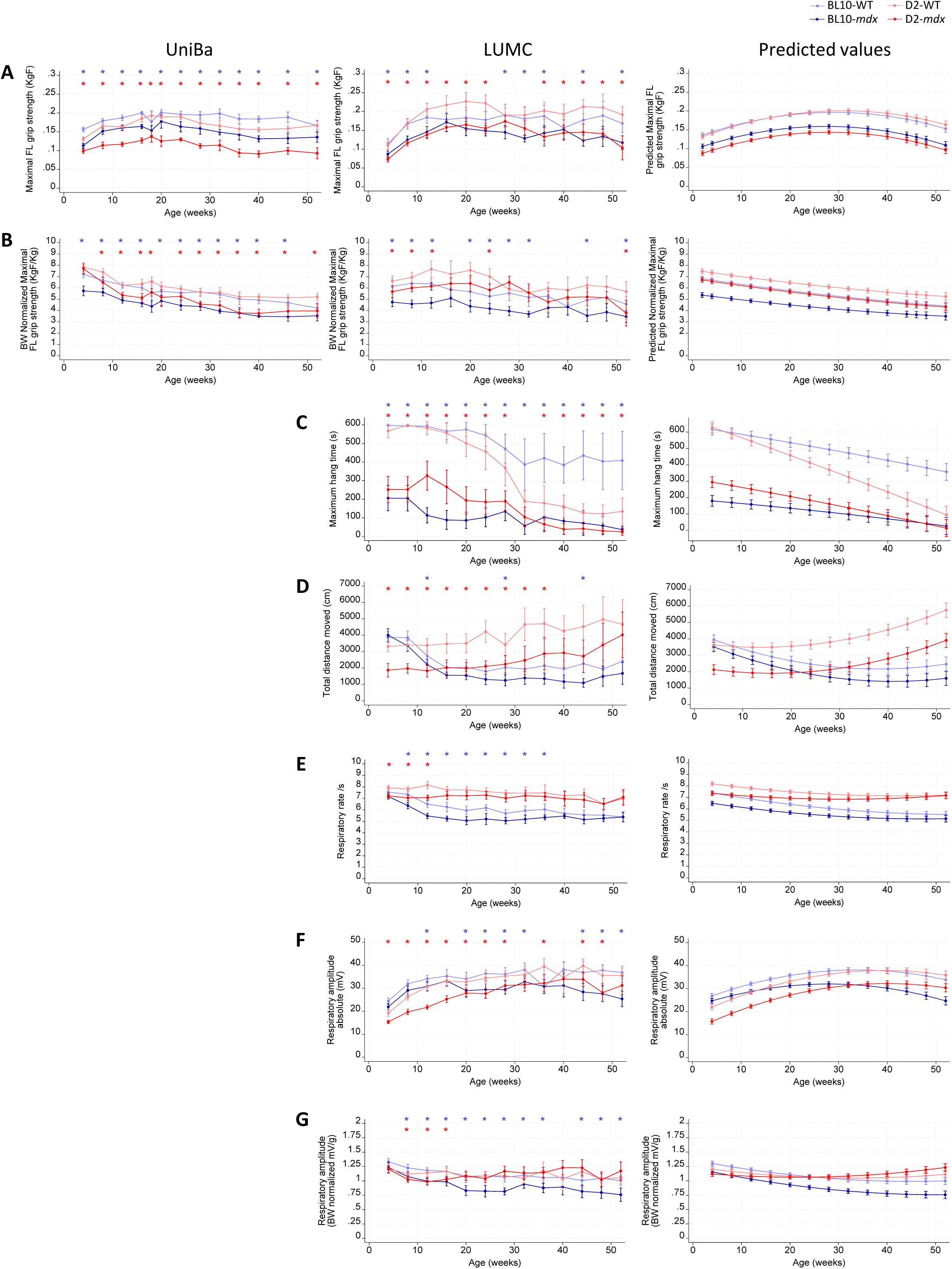
*In vivo* muscle functionality outcome measures. Maximal forelimb grip strength values (A). Body weight normalized maximal grip strength values (B). Absolute and predicted values of the maximal hanging time obtained with the wire hanging test (C) and the total distance moved within 10 minutes in the open field test (D). Whole body plethysmography outcomes including respiratory rate (E), respiratory amplitude (F) and body weight normalized respiratory amplitude (G). Blue and red stars indicate a significant difference (*P*<0.05) between BL10-WT and BL10-*mdx* or D2-WT and D2-*mdx* mice respectively. BW; body weight, FL; forelimb, KgF; kilogram force. UniBa; University of Bari, LUMC: Leiden University Medical Center.

Maximum wire hanging time was largely comparable between the two DMD models, showing a severe impairment that worsened over time (Figure 4C, Supp Table 7). Young adult wildtype mice could hang for the maximum test duration, but their performance declined with age, predominantly in the D2 strain.

General activity was assessed with the open field test. The total distance moved, time moving, and mean velocity were significantly lower in dystrophic mice from both strains versus their wildtypes (Figure 4D, Supp Figure 3 and Supp Table 7). Overall, D2 mice were more active during the test compared to BL10 mice. Furthermore, whereas activity decreased with age in the BL10 strain, it either remained stable or increased in the D2 strain. Dystrophic mice from either strain tended to spend less time in the inner zone than their wildtypes (Supp Figure 3C).

The respiratory rate of both *mdx* models was lower compared to their wildtypes, reaching significance up to 12 (D2-*mdx*) or 36 weeks (BL10-*mdx*) (Figure 4E, Supp Table 8). Notably, D2 animals exhibited higher respiratory rates than BL10 mice during the study period.

Absolute respiratory amplitude was generally lower in both DMD models compared to their wildtypes (Figure 4F). When normalized to BW, respiratory amplitude was lower in BL10-*mdx* mice throughout the study, but only up to the age of 16 weeks in D2-*mdx* mice (Figure 4G). Again, genetic background-related differences were observed: normalized amplitudes of BL10 mice dropped with age, while they remained relatively stable or even increased in D2 animals.

At UniBa, further *in vivo* tests were carried out in mice aged 8, 12, 28, and 52 weeks. Neuromuscular function was assessed by measuring isometric torque of the hindlimb plantar flexor muscles normalized to each mouse BW (Figure 5A-D, Supp Table 9). Torque measurements were carried out longitudinally in all mice cohorts at the main time points, from 8 week onwards, until sacrifice. At all ages, both BL10-*mdx* and D2-*mdx* mice displayed striking lower torque – frequency curves compared BL10-WT and D2-WT, respectively, with a significant reduction in torque at all frequencies of stimulation, from 1 to 200 Hz (Figure 5A-D).

**Figure 5.**
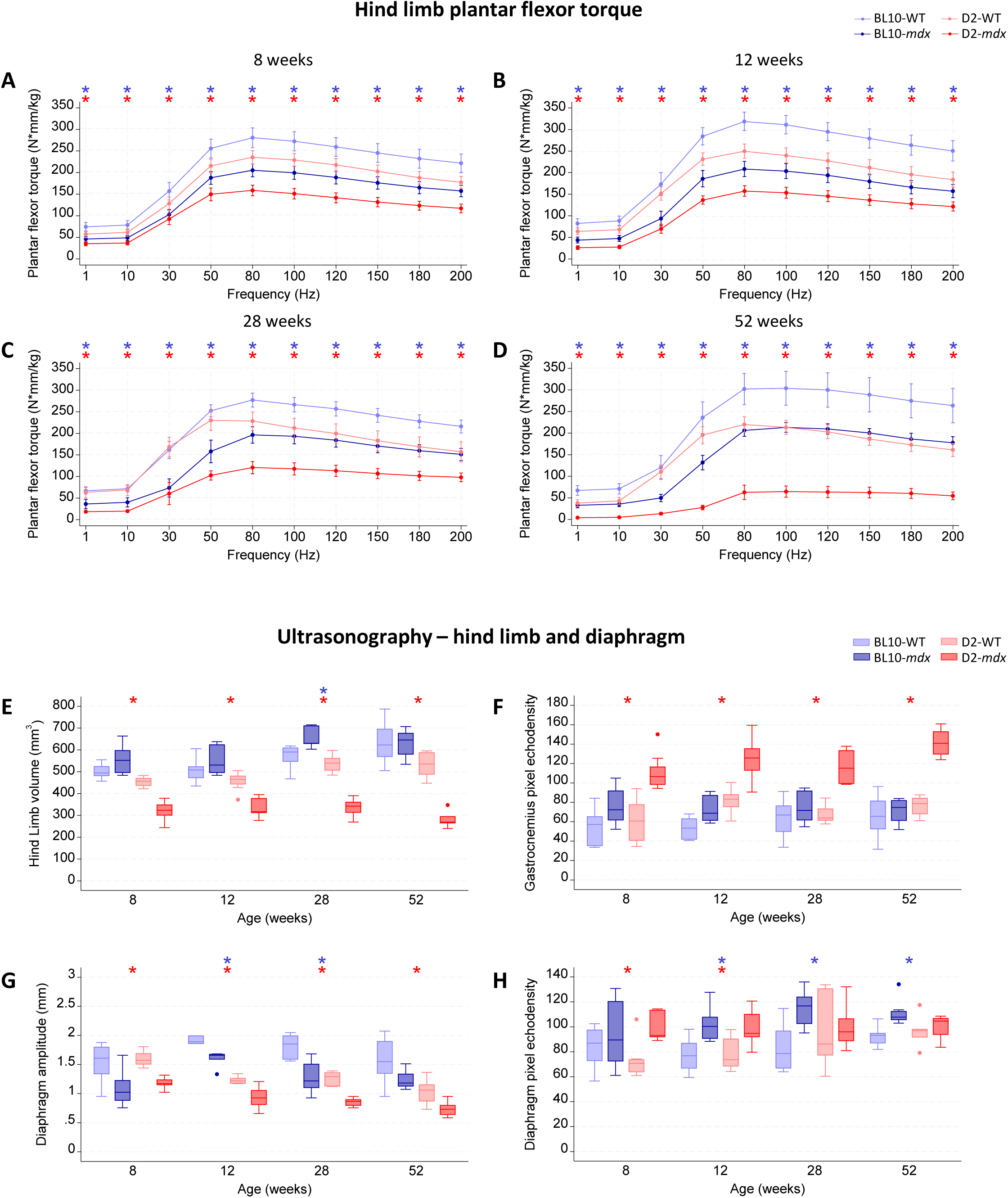
Hindlimb plantar flexor torque and ultrasonography of the hindlimb and diaphragm. Hindlimb plantar flexor torque force frequency curves normalized by body weight acquired at the age of 8 (A), 12 (B), 28 (C) or 52 weeks (D). Body weight normalized hindlimb volume assessed by ultrasonography (E) and mean pixel echodensity of the gastrocnemius (F). Diaphragm movement amplitude (G) and mean pixel echodensity (H) assessed by ultrasonography. Blue and red stars indicate a significant difference (*P*<0.05) between BL10-WT and BL10-*mdx* or D2-WT and D2-*mdx* mice respectively.

Globally, D2-WT mice exhibited lower torque – frequency curves with respect to wildtypes from the BL10 strain, especially at medium – high frequencies of stimulation (≥ 50 – 80 Hz), with differences reaching statistical significance (Supp Table 9). The same trend was observed between dystrophic mice from the two strains, with D2-*mdx* producing significantly lower curves when compared to classic BL10-*mdx* at all ages and frequencies (Supp Table 9). Importantly, while curves recorded from BL10 (WT or *mdx*) and D2-WT mice always remained in about the same range of values over time, a dramatic drop in curves produced by D2-*mdx* mice was observed, particularly at 52 weeks of age (Figure 5D).

In parallel, ultrasound indices of hindlimb and diaphragm muscles were evaluated in mice cohorts at the main endpoints, prior to sacrifice (Figure 5E-H, Supp Table 10). As shown in Figure 5E, BL10-*mdx* mice at 8, 12, and 28 weeks of age displayed a higher volume of the hindlimb compared to age-matched BL10-WT mice, with a significant difference at 28 weeks, reflecting the characteristic pseudo-hypertrophy of this model’s hindlimb muscles (36). This was no longer observed in 52-week-old BL10 mice, most likely due to the stabilization of *mdx* pathology. Differently, D2-*mdx* mice at all ages exhibited a significant decrease in hindlimb volume either compared to their WT (Figure 5E) or to BL10-*mdx* mice (Supp Table 10), in line with the atrophic phenotype observed in this model. Overall, healthy D2 mice also showed smaller hind limb volumes compared with BL10 wildtype mice (Supp Table 10). Additionally, gastrocnemius muscles of BL10-*mdx* tended to show increased echodensity compared to their wildtypes, particularly in the acute phase of pathology at 8 and 12 weeks of age (Supp Table 10). Interestingly – coherently with their reported hyperfibrotic disease profile (18) – D2-*mdx* mice exhibited a significant increase in gastrocnemius echodensity as early as 8 weeks of age and continuing up to 52 weeks (Figure 5F). D2-WT values were largely comparable with those obtained in BL10 controls over time, except for a significant increase in echodensity at 12 weeks of age (Supp Table 10).

Focusing on the diaphragm, respiratory movement amplitude was severely compromised in *mdx* mice of both strains compared with their respective controls. In BL10-*mdx* mice, these differences reached statistical significance only at 12 and 28 weeks of age, whereas in D2-*mdx* significance was observed at all ages (Figure 5G, Supp Table 10). Interestingly, while inter-strain values nearly overlapped at 8 weeks of age, from 12 weeks onwards a significant decrease in amplitude was observed in D2 mice, both WT and *mdx*, compared with their BL10 counterparts (Supp Table 10). Diaphragm echodensity (Figure 5H) was significantly higher in BL10-*mdx* mice versus their wildtypes from 12 weeks of age onwards. In contrast, in D2-*mdx* mice, this increase relative to D2-WT mice was already evident at 8 weeks of age and remained present at 12 weeks, but not at later time points. No significant differences were observed between BL10 and D2 mice (Supp Table 10).

### 3.3 D2-mdx mice develop cardiomyopathy at an earlier age than BL10-mdx mice

Left ventricular function was assessed prior to sacrifice at the age of 8, 12, 28 and 52 weeks using either ultrasonography (UniBa) or magnetic resonance imaging (MRI; LUMC) (Figure 6, Supp Table 11). Overall, no alterations were observed in mice aged 8 or 12 weeks regardless of the strain. Stroke volume was decreased in D2-*mdx* mice from 28 weeks of age onwards, whereas BL10-*mdx* mice developed this reduction later at 52 weeks of age (Figure 6A). Alterations in cardiac output, ejection fraction and fractional shortening showed an identical pattern (Figure 6B-D). Values of these indices were reduced in *mdx* mice of both strains compared to their wildtypes, reaching significance from the age of 28 weeks in D2-*mdx* mice, and from 52 weeks in BL10-*mdx* mice. Furthermore, it should be pointed out that also D2-WT mice showed an impairment in ejection and shortening fractions, with significant reductions compared to BL10-WT mice at the age of 52 weeks.

**Figure 6.**
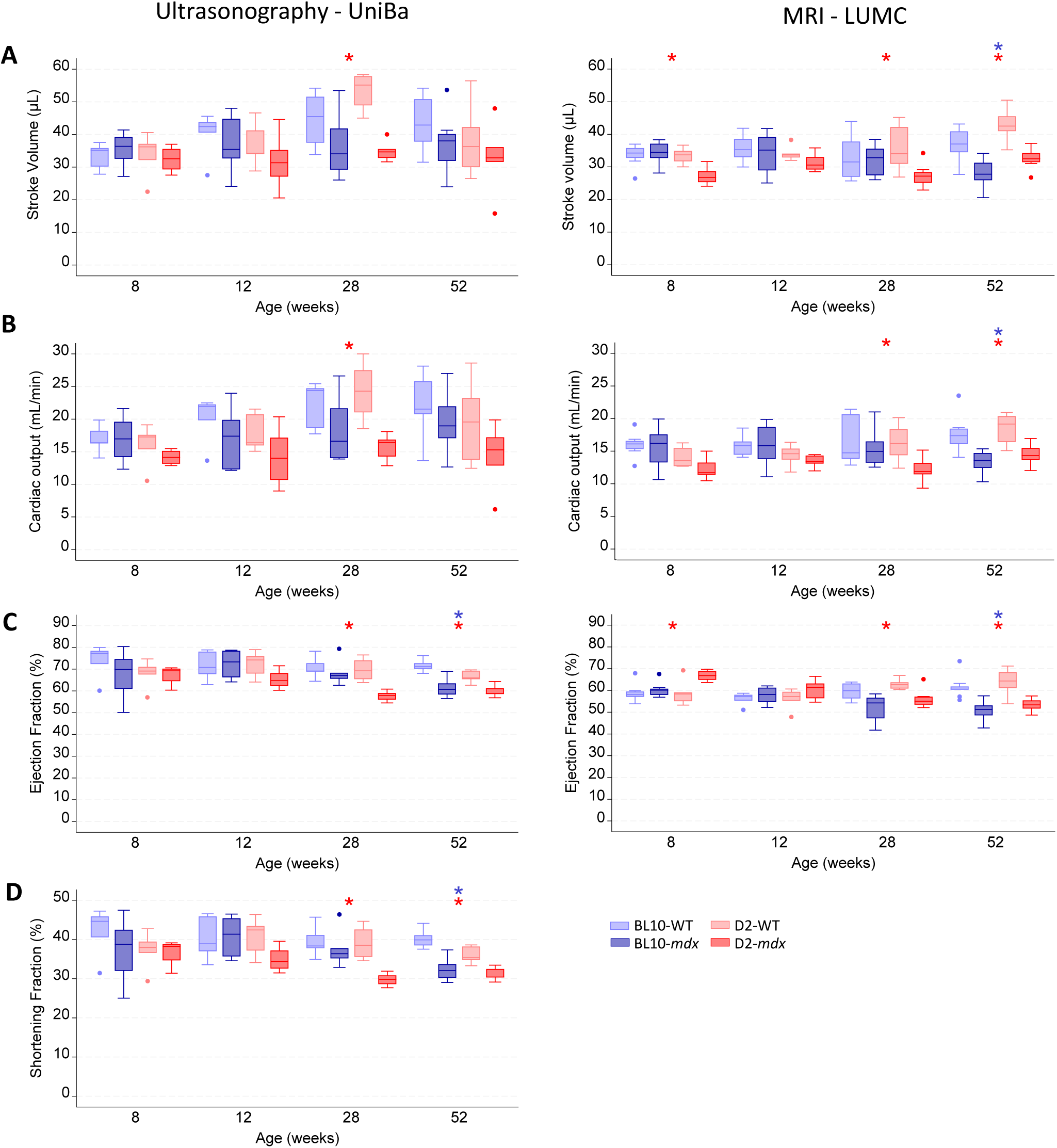
Heart functionality. Left ventricular function was assessed by echocardiography (UniBa) or magnetic resonance imaging (LUMC) at the endpoint. Stroke volume (A), cardiac output (B), ejection fraction (C) and shortening fraction (D) were assessed from mice aged 8 weeks or older. Blue and red stars indicate a significant difference (*P*<0.05) between BL10-WT and BL10-*mdx* or D2-WT and D2-*mdx* mice respectively. UniBa; University of Bari, LUMC: Leiden University Medical Center.

### 3.4 Isolated muscle contractile function is similarly affected in both dystrophic mouse models

As the final step in characterizing D2-*mdx* versus BL10-*mdx* mice, further assessments at UniBa were aimed at measuring isometric and eccentric contraction parameters in freshly isolated EDL and diaphragm muscles from each experimental cohort at 8, 12, 28, and 52 weeks of age.

Data for EDL muscle isometric and eccentric contraction are shown in Figure 7, Supp Figure 5, and Supp Table 12. With respect to single twitch kinetics, values for both TTP and HRT showed modest fluctuations across mice groups at every age (Supp Figure 5A-B, Supp Table 12). Specific twitch and tetanic force values were consistently lower in either BL10 or D2 dystrophic mice compared with their wildtype counterparts (Figure 7A-B); this difference was statistically significant at all ages, except for D2-*mdx* mice at 52 weeks. Inter-genotype differences were not significant in WT or *mdx* mice, despite a trend toward lower values in D2-WT mice observed from 12 weeks onwards (Supp Table 12). For completeness, absolute force values were also included (Supp Figure 5C-D). Although generally resembling the differences observed for normalized data, in this case, D2-*mdx* mice appeared significantly weaker than BL10-*mdx* (Supp Table 12). This discrepancy reflects the difference in EDL muscle weight between the two models (Supp Figure 1), which is not considered when measuring absolute force. Upon eccentric stimulation, EDL muscles from BL10-*mdx* mice showed a significant force drop versus BL10-WT mice at the 5^th^ and 10^th^ pulses (Figure 7C-D) in 28- and 52-week-old animals; D2-*mdx* mice suffered the most at the 10^th^ pulse in the same age windows (Figure 7D). Dystrophic EDL muscles showed little recovery from the stretch-induced force loss after either a 5- or 15-minute pause, irrespective of the genotype (Figure 7E-F). A comparable behavior in response to stretch was observed in either BL10 or D2-WT animals (Supp Table 12).

**Figure 7.**
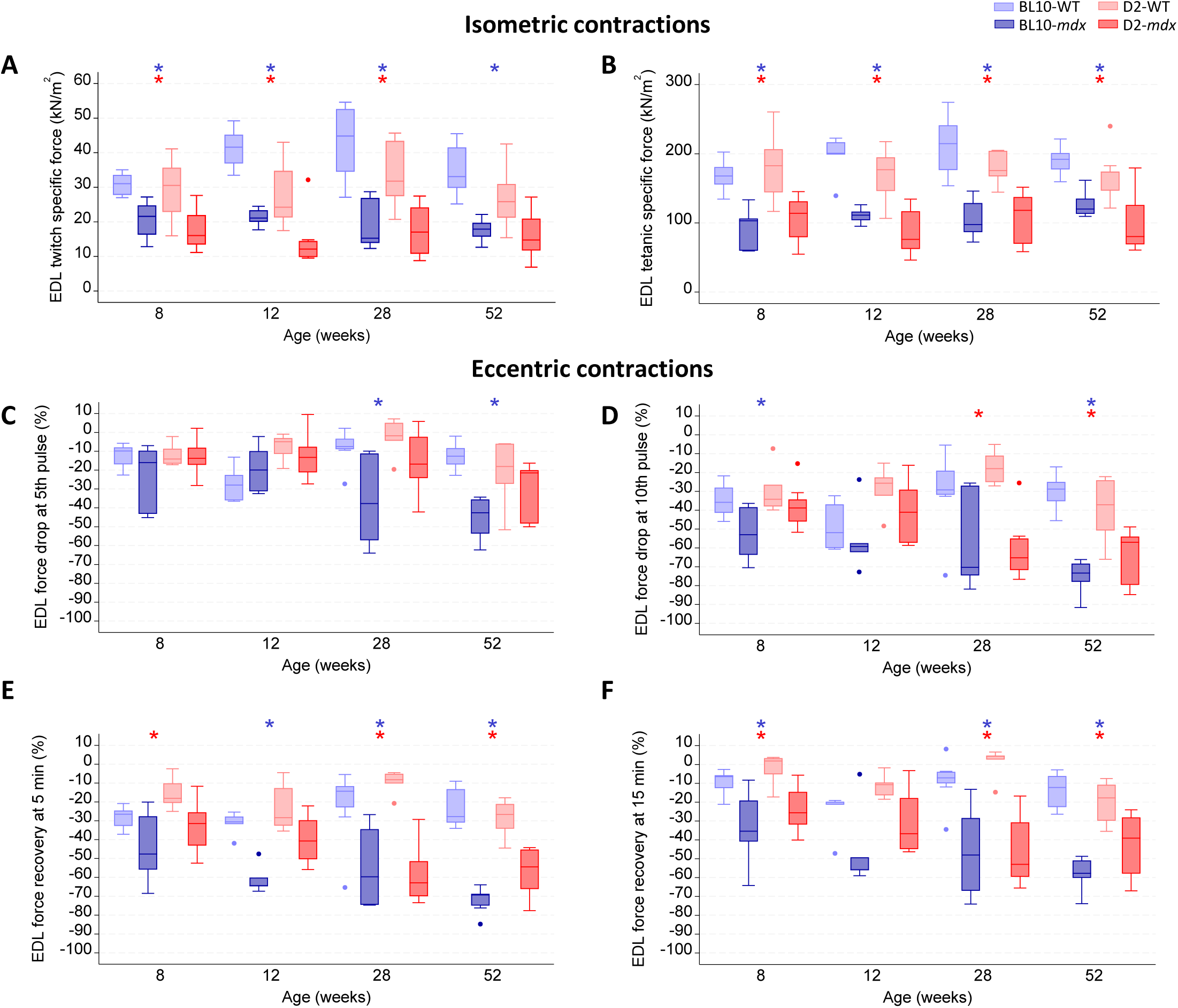
*Ex vivo* physiology assessments of the extensor digitorum longus. Isometric contractions consisting of twitch (A) and tetanic (B) specific forces were determined first, followed by 10 consecutive eccentric contractions (10% lengthening) of which the force drop from baseline (first pulse) is shown for the 5^th^ contraction (C) and the 10^th^ contraction (D). Force recovery was assessed at 5 min (E) and 15 min (F) after the eccentric contraction protocol. Blue and red stars indicate a significant difference (*P*<0.05) between BL10-WT and BL10-*mdx* or D2-WT and D2-*mdx* mice respectively. EDL; extensor digitorum longus.

Data for diaphragm isometric and eccentric contraction are shown in Figure 8, Supp Figure 6, and Supp Table 13. Single twitch kinetics showed only modest differences in TTP and HRT across cohorts, except for a significant increase in HRT in 52-week-old BL10-*mdx* mice versus wildtype mice (Supp Figure 6A–B, Supp Table 13). For diaphragm, absolute force values are not meaningful due to the nature of the *ex vivo* preparation; therefore, only specific force values were reported herein. Twitch and tetanic force were consistently reduced in dystrophic mice of both strains versus wildtypes, reaching significance from the age of 12 weeks (Figure 8A-B, Supp Table 13). Although no significant intergenotype differences were found, D2-WT mice tended to show lower force values compared to BL10 healthy controls. In response to stretch, 12-week and 28-week-old BL10-*mdx* mice showed a significant force drop at the 5^th^ or 10^th^ pulse, while for D2-*mdx* this was observed at 28 and 52 weeks of age (Figure 8C-D). Similarly, dystrophic mice from both strains were mostly unable to recover from the eccentric protocol at all ages, except for 8 weeks (Figure 8E-F). Again, no inter-genotype differences were found on these indices (Supp Table 13).

**Figure 8.**
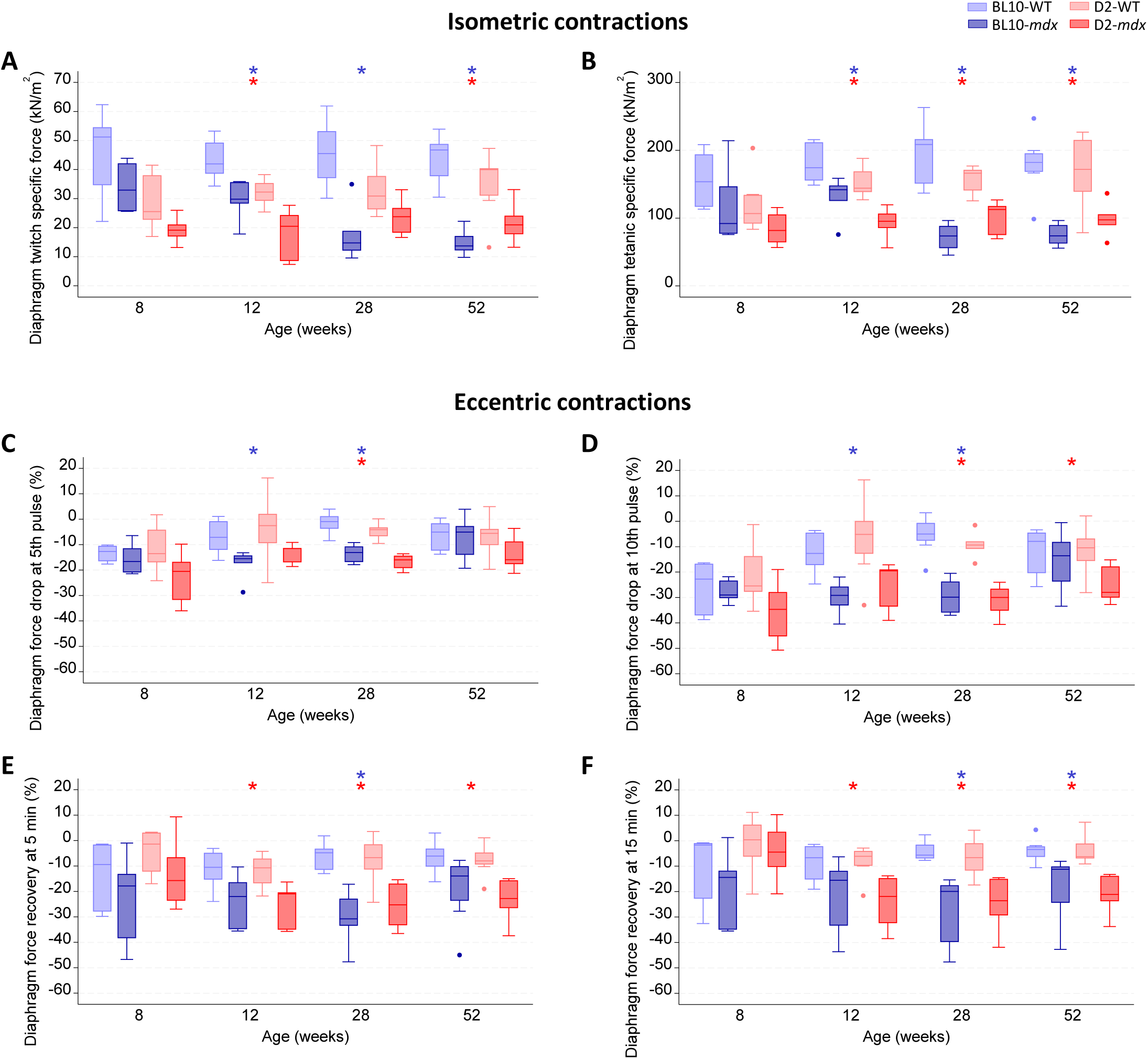
*Ex vivo* physiology assessments of the diaphragm. Isometric contractions consisting of twitch (A) and tetanic (B) specific forces were determined first. Directly thereafter, 10 consecutive eccentric contractions involving 10% muscle lengthening were performed. The force reduction from baseline (first pulse) for the 5^th^ (C) and 10^th^ (D) contractions were recorded. At 5 (E) and 10 (F) minutes after the eccentric contraction protocol’s completion, force recovery was assessed. Blue and red stars indicate a significant difference (*P*<0.05) between BL10-WT and BL10-*mdx* or D2-WT and D2-*mdx* mice respectively.

### 3.5 Plasma CK and LDH levels are elevated in both mdx models, but drop in D2-mdx with age

To gain insight into muscle integrity throughout life, CK and LDH levels were measured in plasma samples obtained from mice at terminal time points (Figure 9 and Supp Table 14). Although absolute CK values differed between the study sites, likely due to methodological differences, similar patterns were identified (Figure 9A). CK levels were low and did not differ between the WT strains. BL10-*mdx* mice had elevated plasma CK levels compared to BL10-WT mice already from 2-weeks-of-age onwards. In line, CK levels of D2-*mdx* mice exceeded those of D2-WT mice between the age of 8-28 weeks. Notably, aged D2-*mdx* mice had lower CK levels than age-matched BL10-*mdx* mice which could relate to their muscle atrophy.

**Figure 9.**
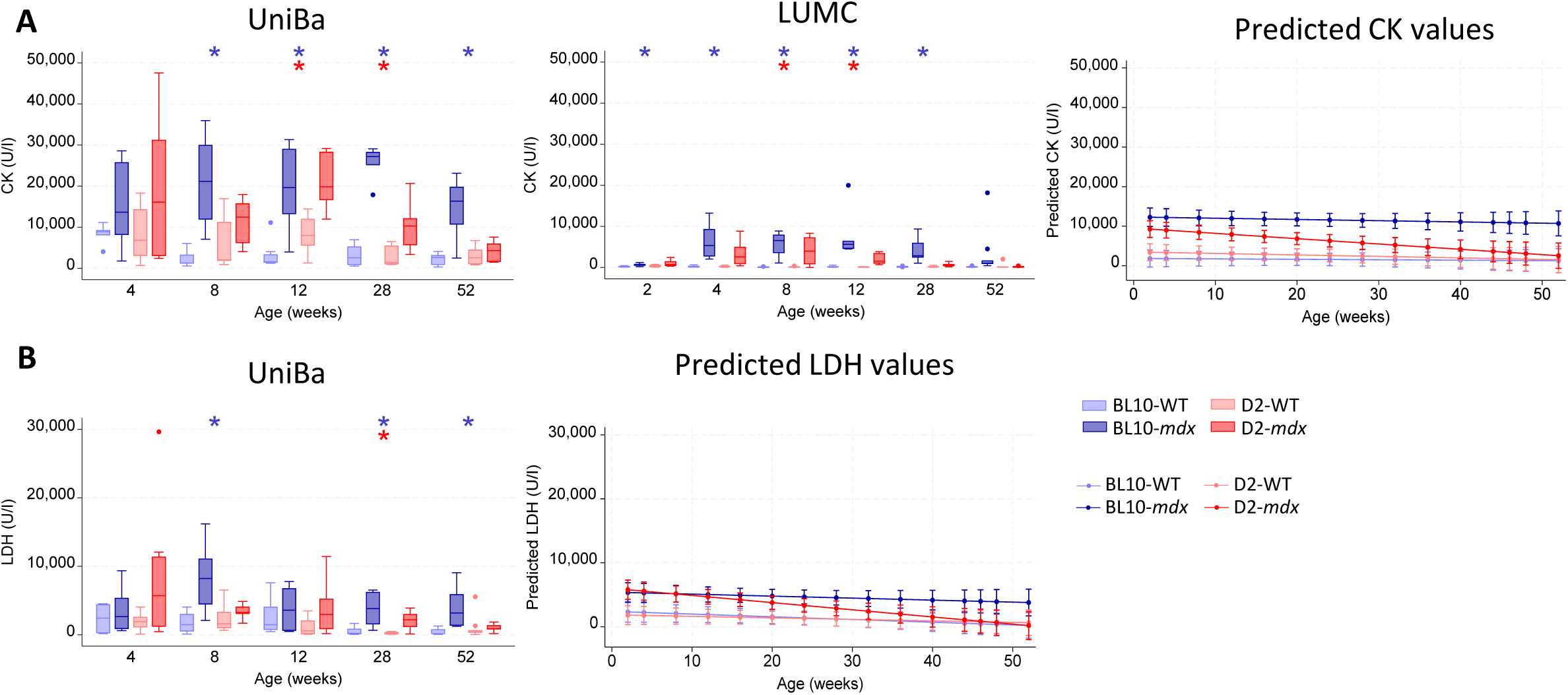
Plasma creatine kinase and lactate dehydrogenase levels. Prior to sacrifice, CK (A) and LDH (B) levels were assessed. The absolute and predicted values are shown. Blue and red stars indicate a significant difference (*P*<0.05) between BL10-WT and BL10-*mdx* or D2-WT and D2-*mdx* mice respectively. CK; creatine kinase, LDH; lactate dehydrogenase. UniBa; University of Bari, LUMC: Leiden University Medical Center.

Plasma LDH levels were increased in both DMD models compared to healthy wildtype mice, reaching significance predominantly in the BL10-*mdx* mice (Figure 9B). In line with CK levels, LDH levels also decreased in D2-*mdx* with age, but not in BL10-*mdx* mice.

## 4. Discussion

The D2-*mdx* mouse has gained attention from the DMD research community because of its more severe disease phenotype compared to the classic BL10-*mdx* mouse. In recent years, an increasing number of studies have been conducted using the D2-*mdx* model, despite the lack of a thorough and comprehensive characterization of its disease course. In light of this, the OMAM initiative was conceived with the overall aim to provide guidance on the most appropriate use of the D2-*mdx* mouse model in preclinical research. A preliminary step involving an extensive revision and collective analysis of available data was performed to identify knowledge gaps in this model. This revealed that the results of previous studies were relatively fragmented and that there were methodological differences between groups on several outcome measures (37). The follow-up study, presented here, was then dedicated to extensively characterize the natural history of the D2-*mdx* mouse, compared to classic BL10-*mdx*, by two independent laboratories. In this regard, despite the inherent challenges of a multi-site design, this approach provided complementary and robust insights, increasing result reliability and enabling cross-validation and pooled analyses to enhance statistical power.

Overall, data obtained by the two laboratories were largely comparable, highlighting the suitability of the existing TREAT-NMD SOPs for the analysis of disease-related readouts in both models. One key distinguishing feature when comparing these strains is body composition, with muscle pseudohypertrophy observed in BL10-*mdx* mice (33–35) and atrophy in D2-*mdx* mice(38). Indeed, BL10-*mdx* mice generally had a larger body, and larger lean and muscle masses compared to D2-*mdx* mice. The pronounced muscle atrophy in D2-*mdx* mice was also highlighted by ultrasound data on hindlimb volume at all ages. From the analysis of body composition, it emerged that the amount of fat was comparable between D2-*mdx* and D2-WT mice, while BL10-*mdx* mice exhibited lower fat mass. We speculate that, in this case, fat tissue was largely consumed to accommodate for maintenance of BL10 dystrophic and hypertrophic muscles (39,40). At the muscle-specific level, the hyper-fibrotic nature of the D2-*mdx* model could account for the significant increase in gastrocnemius ultrasound echodensity observed throughout the study. Notably, both wildtype and dystrophic D2 mice, had a lower body and muscle weight compared to the BL10 background, which should be acknowledged when testing drugs targeting muscle atrophy. The D2-*mdx* mice atrophic phenotype could also account for the reduction in plasma CK and LDH levels observed with age, which was not found in BL10-*mdx* mice.

Muscle functionality was investigated with a panel of disease relevant assessments, both *in vivo* and *ex vivo*. *In vivo* results from both sites were consistent in terms of absolute forelimb force, with D2-*mdx* mice performing significantly worse than the other cohorts, including BL10-*mdx*. However, this trend became less evident after normalization to body weight, particularly when considering predicted values, where classic *mdx* mice emerged as the weakest cohort. While grip strength remains a valid outcome measure for both models since the dystrophy-related weakness is clearly appreciable, data normalization may need to account for additional factors, such as differences in body composition between the two strains, to more accurately reflect functional deficits.

*In vivo* neuromuscular function (*i.e.,* plantar flexor torque), was consistently impaired in D2-*mdx* mice compared with all other cohorts, and this deficit worsened over time. Importantly, D2 mice – including wildtypes – produced much lower forces than their BL10 counterparts, indicating an intrinsic weakness of this strain that remained evident even after normalization to body weight. This strong influence of genetic background on this parameter should therefore be carefully considered when using it in this model.

Indeed, for other outcome measures, such as the open field test, differences between genetic backgrounds were similarly observed and even exceeded the variations between dystrophic and wildtype mice within each strain. Notably, D2 mice exhibited higher locomotor activity in the open field, likely reflecting strain-specific behavioral traits rather than disease-related effects. These observations underscore the strong influence of genetic background on certain parameters, highlighting the need for strain-matched wild-type controls and caution in interpreting the D2-*mdx* phenotype as uniformly more severe across all outcome measures.

*Ex vivo* muscle physiology of either EDL or diaphragm, was equally affected by the pathology in both models, irrespective of genetic background. Nonetheless, even in this case, including strain-matched controls remains crucial given the observed differences in hind limb muscle weight and to ensure experimental rigor.

For the analysis of cardiac function, two complementary techniques were used (ultrasonography and MRI), providing comparable outcomes. Both DMD models displayed a reduction in ejection fraction, stroke volume and – for echocardiography – shortening fraction, compared to their wildtypes. Notably, reduced ejection and shortening fractions manifested earlier in D2-*mdx* mice than in BL10-*mdx* mice, confirming a more precocious onset of left ventricular dysfunction in this model (11,41–43). In D2-*mdx* mice, an earlier disease manifestation was also observed at the respiratory level: diaphragm ultrasonography, in line with whole-body plethysmography findings, revealed reduced respiratory amplitude compared with classic *mdx* mice, already detectable at 12 weeks of age. These results underline the importance of considering the timing of disease-related alterations when designing preclinical studies aiming at the pharmacological targeting of cardiac and respiratory aspects of DMD. Furthermore, we also confirmed previous studies highlighting that the heart weight of D2-*mdx* mice is greater than that of BL10-*mdx* mice; this may be, at least in part, due to the presence of calcifications previously described in this model and not found in classic *mdx* (20,44).

## 5. Conclusions

In conclusion, this study provides a comprehensive overview of the natural disease history of D2-*mdx* and BL10-*mdx* mice, highlighting the timing and progression of key pathological events. The high degree of consistency across independent laboratories supports the validity of TREAT-NMD SOPs and reinforces the importance of standardization and reproducibility in preclinical research. Predicted values derived from these data could represent a resource for designing future studies, enhancing both research quality and reproducibility. Moreover, the observed differences between strains offer guidance for selecting the most appropriate model based on the specific pathological mechanisms and outcome measures under investigation. In particular, our findings suggest that, given its characteristics, the D2-*mdx* model could be especially useful for evaluating potential interventions aimed at counteracting skeletal muscle atrophy or slowing the progression of cardiomyopathy. At the same time, we reiterate the need for caution when interpreting readouts for which intrinsic deficits are already present in wildtype D2 mice. Further analyses dedicated to the assessment of molecular and histological features of skeletal, respiratory, and cardiac muscles in the same experimental groups will complement the functional findings presented here, potentially enabling correlations across multiple readouts, as previously demonstrated within the OMAM initiative (45).

## Acknowledgements

This study has been funded by Charley’s Fund and Duchenne UK, in the framework of the OMAM “Of Mice And Measures” project, with funds granted to Annemieke Aartsma-Rus and Annamaria De Luca.

## List of abbreviations

DMD: Duchenne muscular dystrophy
BL10-*mdx*: C57BL/10ScSn-*Dmd^mdx^*/J mouse
D2-*mdx*: D2.B10-*Dmd^mdx^*/J mouse
BW: Body weight
WT: Wildtype
DGC: Dystrophin-glycoprotein complex
TTP: Time to peak
HRT: Half-relaxation time
LUMC: Leiden University Medical Center
UniBa: University of Bari Aldo Moro
MRI: Magnetic resonance imaging
EDL: Extensor digitorum longus
EF: Ejection fraction
SF: Shortening fraction
SV: Stroke volume
CO: Cardiac output
CK: Creatine kinase
LDH: Lactate dehydrogenase
L□: Optimal length
ESV: End-systolic volume
EDV: End-diastolic volume
HR: Heart rate
PLAX: Parasternal long axis
LV: Left ventricle
SOL: Soleus
LTBP4: Latent TGF-β-binding protein 4
Dmd: Gene encoding dystrophin
OMAM: “Of Mice and Measures” project

## Supplementary Figures

**Supp Figure 1. Absolute muscle weights.** Absolute muscle weights and corresponding predicted values for the quadriceps (A), gastrocnemius (B), tibialis anterior (C), triceps (D), heart (E) and EDL (F). Blue and red stars indicate a significant difference (*P*<0.05) between BL10-WT and BL10-*mdx* or D2-WT and D2-*mdx* mice respectively. EDL; extensor digitorum longus. UniBa; University of Bari, LUMC: Leiden University Medical Center.

**Supp Figure 2. Organ weights and bone length.** Absolute and body weight normalized organ weights and corresponding predicted weight values for the kidney (A), liver (B) and spleen (C). Length assessments of the tibia (D) and femur (E). Blue and red stars indicate a significant difference (*P*<0.05) between BL10-WT and BL10-*mdx* or D2-WT and D2-*mdx* mice respectively. UniBa; University of Bari, LUMC: Leiden University Medical Center.

**Supp Figure 3. Spontaneous activity.** Absolute and predicted values of the total time moving (A), mean velocity (B) and time spent in the inner zone of the box (C), assessed with the open field test on a monthly basis. Blue and red stars indicate a significant difference (*P*<0.05) between BL10-WT and BL10-*mdx* or D2-WT and D2-*mdx* mice respectively.

**Supp Figure 4. Predicted values of heart functionality.** Predicted values for the left ventricular stroke volume (A), cardiac output (B), ejection fraction (C) and shortening fraction (D), assessed either by ultrasonography (left panels) or magnetic resonance imaging (right panels). UniBa; University of Bari, LUMC: Leiden University Medical Center.

**Supp Figure 5. *Ex vivo* physiology of the extensor digitorum longus.** Time to peak (A), half-relaxation time (B) and maximal force (C) of twitch contractions, and tetanic maximal force (D) shown as absolute and predicted values. Predicted values of the specific force evoked by twitch (E) and tetanic (F) contractions. Predicted values of the force drop at the 5^th^ (G) and 10^th^ (H) contraction in a series of 10 eccentric contractions compared to the baseline contraction. Predicted force recovery of the EDL at 5 (I) and 10 minutes (J) after the eccentric contraction protocol. Blue and red stars indicate a significant difference (*P*<0.05) between BL10-WT and BL10-*mdx* or D2-WT and D2-*mdx* mice respectively. EDL; extensor digitorum longus, TTP; time to peak, HRT; half-relaxation time.

**Supp Figure 6. *Ex vivo* physiology of the diaphragm.** Time to peak (A), half-relaxation time (B), and the specific force (C) were analyzed of twitch contractions and reported as absolute and predicted values. Predicted values for specific force generated by twitch (E) and tetanic (F) contractions were calculated. Additionally, predicted force reductions at the 5^th^ (G) and 10^th^ (H) eccentric contraction during a series of 10 eccentric contractions were compared to baseline. Finally, predicted recovery of EDL force was assessed at 5 (I) and 10 minutes (J) following completion of the eccentric contraction protocol. The blue star indicates a significant difference (*P*<0.05) between BL10-WT and BL10-*mdx* mice. TTP; time to peak, HRT; half-relaxation time.

