## Supplementary figures and images for "The D2-*mdx* mouse as a preclinical model for Duchenne muscular dystrophy: a natural history study across two independent sites"

### Supplementary Figure 1

#### Supplementary Figure 1

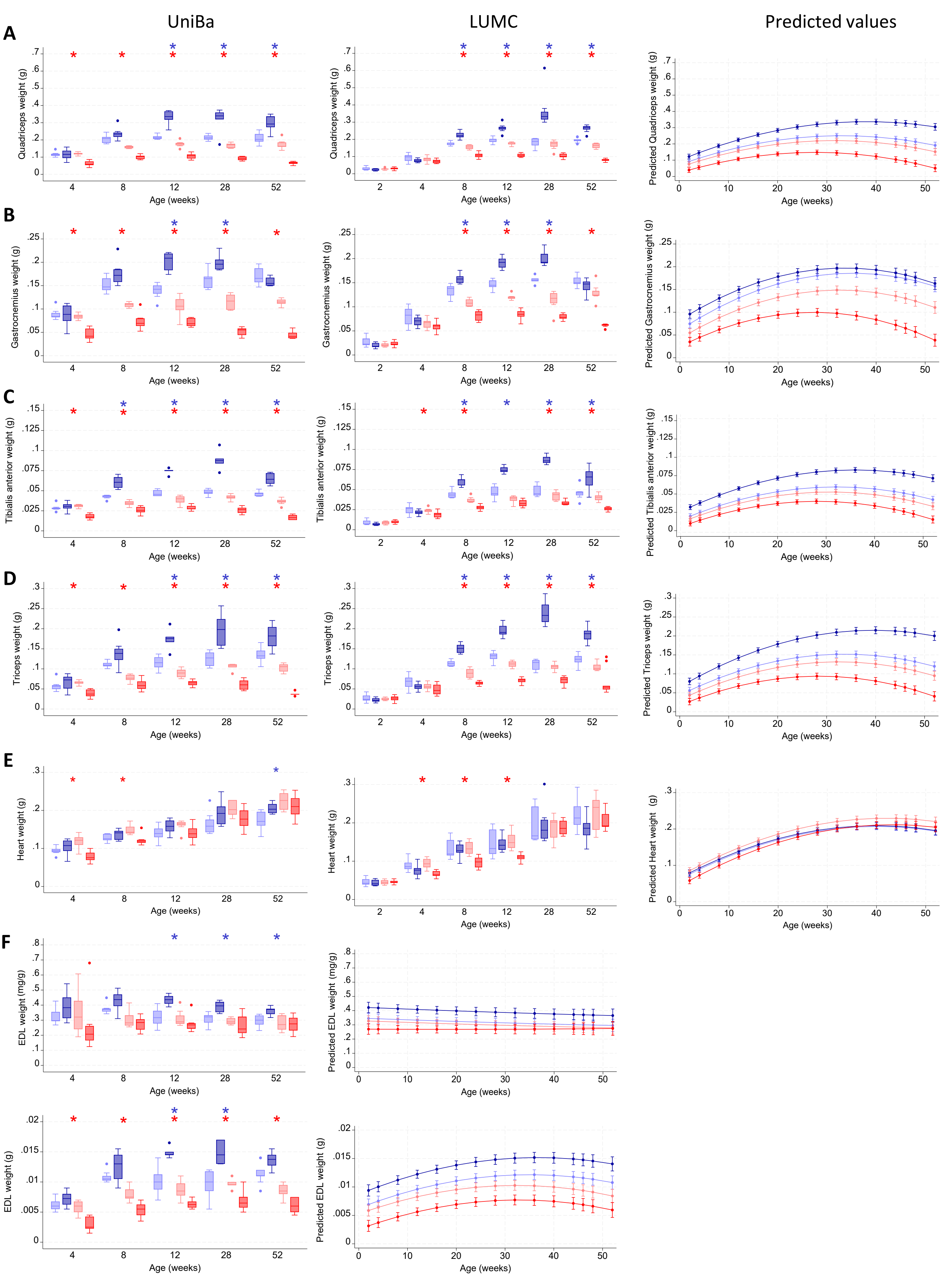

### Supplementary Figure 2

#### Supplementary Figure 2

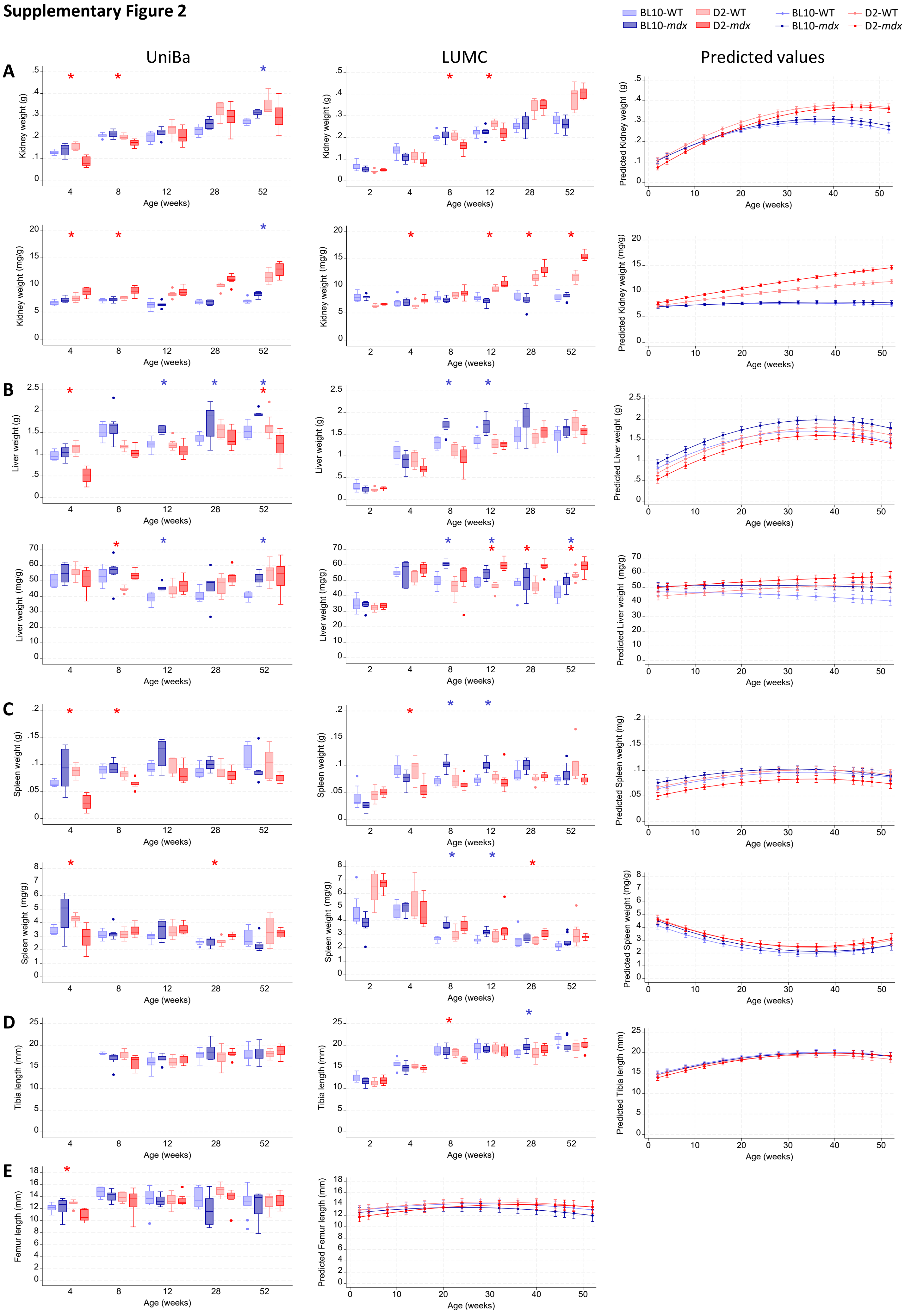

### Supplementary Figure 3

**Supplementary figure 3**

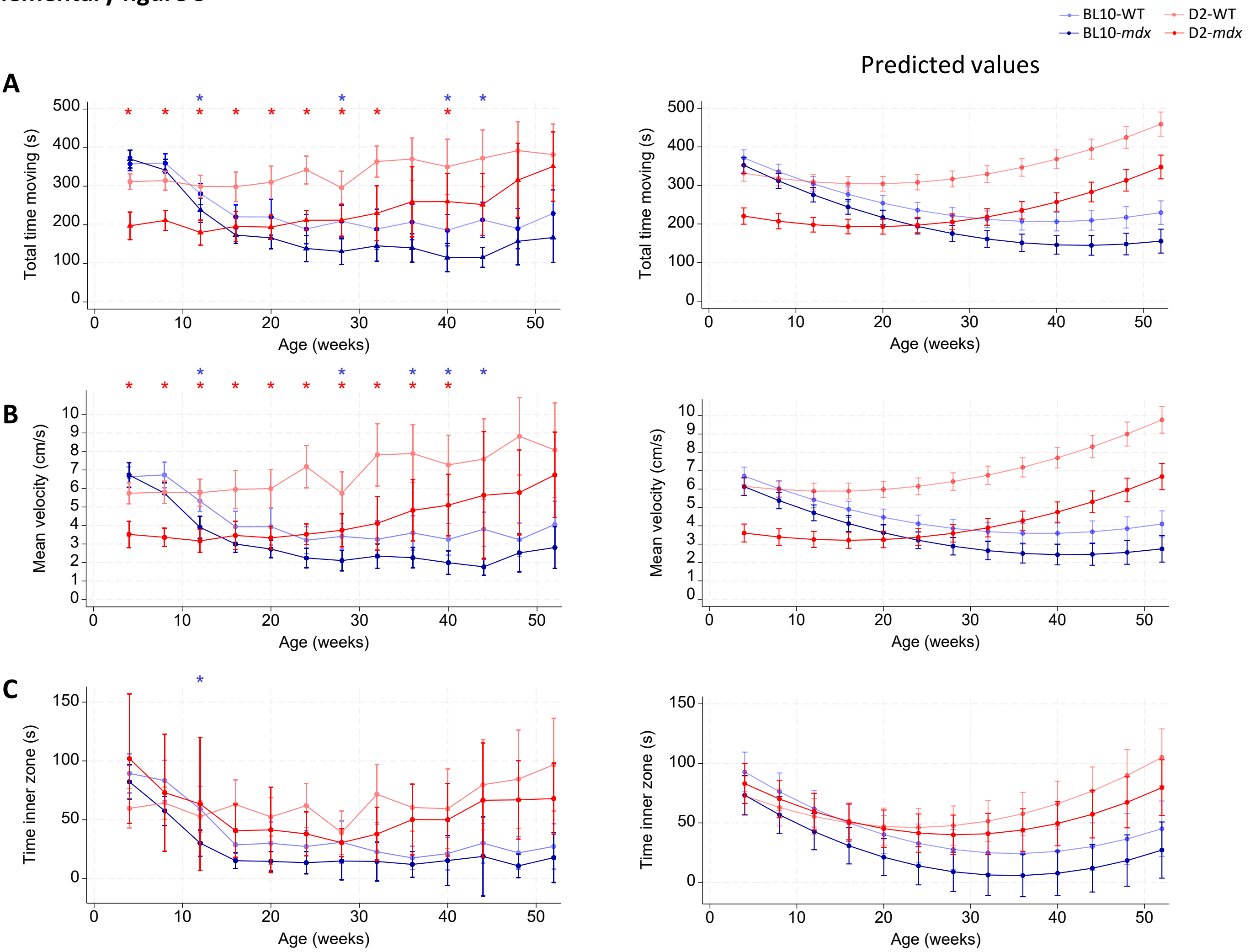

### Supplementary Figure 4

**Supplementary figure 4**

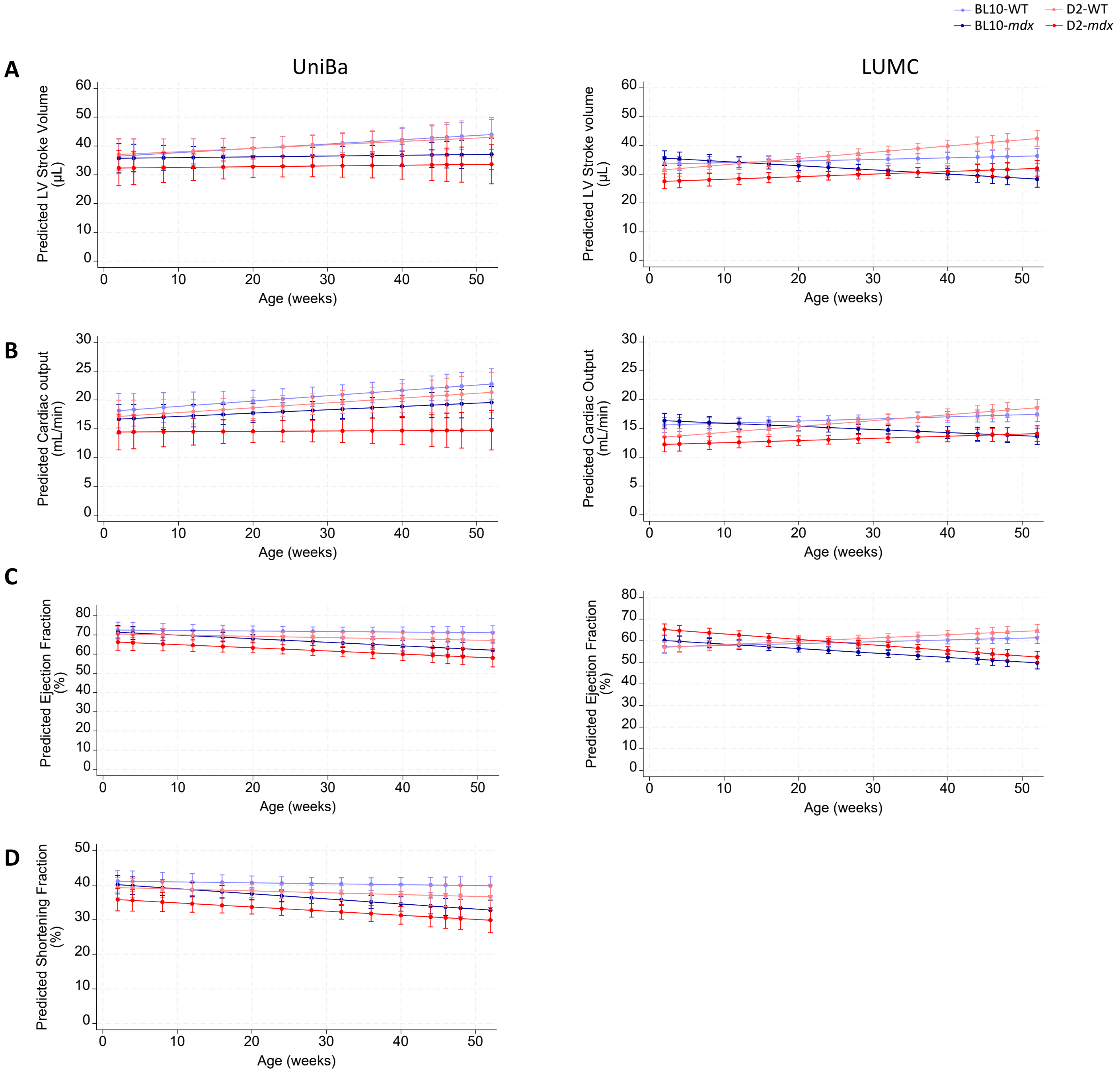

### Supplementary Figure 5

**Supplementary figure 5**

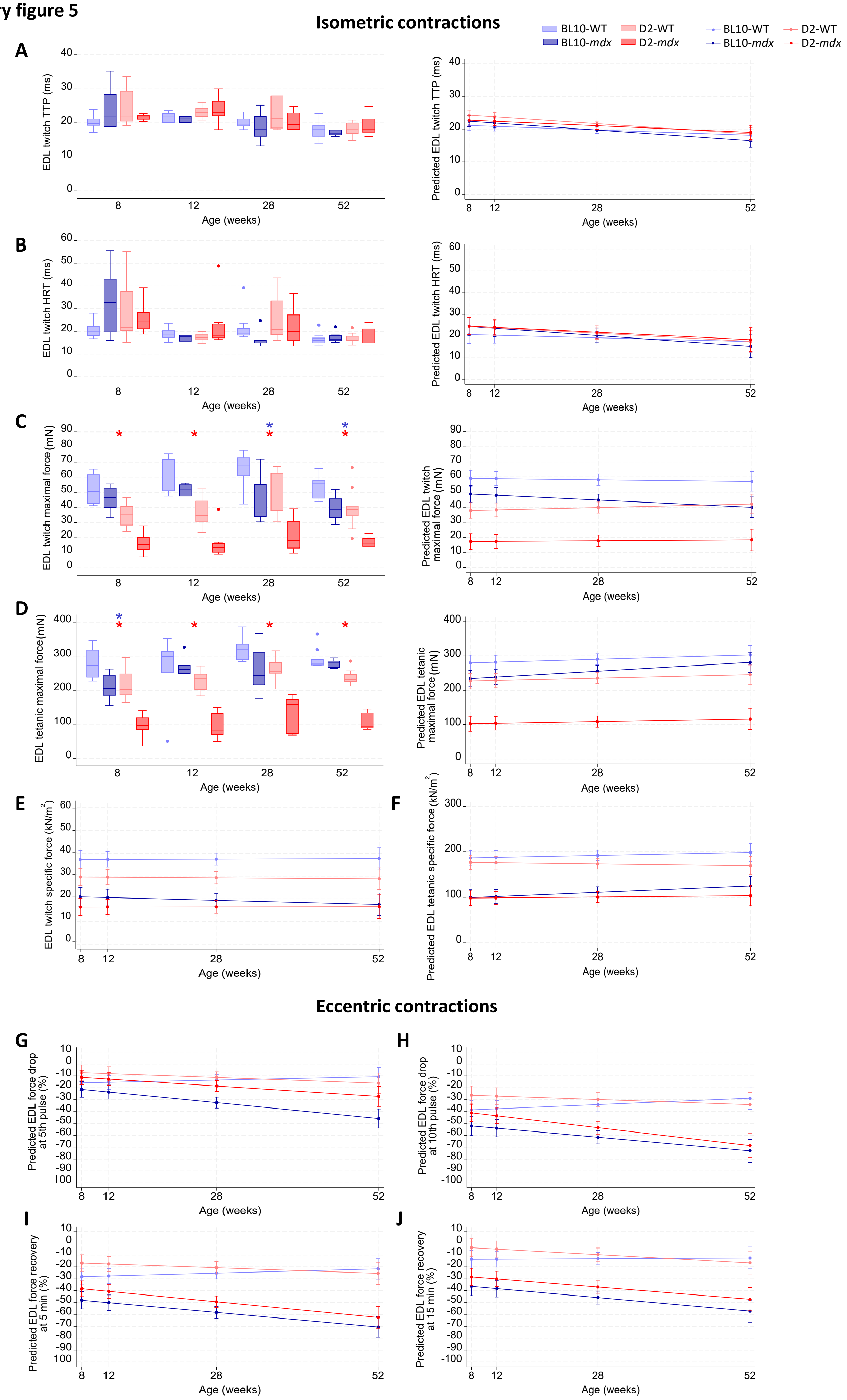

### Supplementary Figure 6

**Supplementary figure 6**

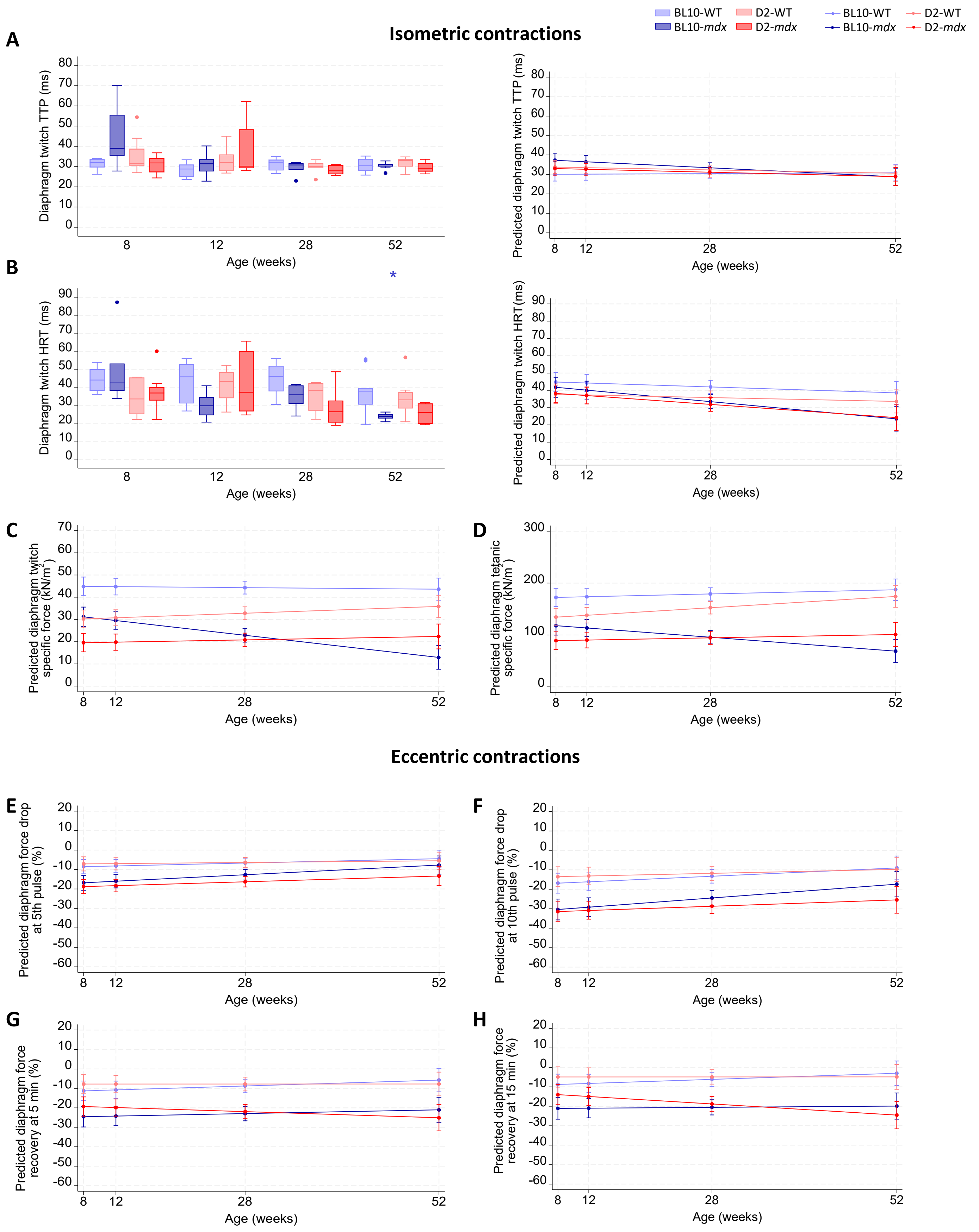
