## Supplementary Tables for "The D2-*mdx* mouse as a preclinical model for Duchenne muscular dystrophy: a natural history study across two independent sites"

| **Outcome measure** | **Coefficient** | ***P*-value** | **Direction** |
| --- | --- | --- | --- |
| Body weight | 1.601 | **0.034** | UniBa > LUMC |
| Quadriceps weight | 0.031 | **0.001** | UniBa > LUMC |
| Gastrocnemius weight | 0.030 | **<0.001** | UniBa > LUMC |
| Tibialis anterior weight | -0.022 | **<0.001** | UniBa < LUMC |
| Triceps weight | 0.017 | **<0.001** | UniBa > LUMC |
| Heart weight | 0.021 | **<0.001** | UniBa > LUMC |
| Kidney weight | 0.018 | 0.08 |  |
| Liver weight | 0.158 | **0.002** | UniBa > LUMC |
| Spleen weight | 0.012 | **<0.001** | UniBa > LUMC |
| BW normalized quadriceps weight | 0.509 | **0.013** | UniBa > LUMC |
| BW normalized gastrocnemius weight | 0.467 | **0.001** | UniBa > LUMC |
| BW normalized tibialis anterior weight | -0.872 | **<0.001** | UniBa < LUMC |
| BW normalized triceps weight | 0.002 | 0.99 |  |
| BW normalized heart weight | 0.044 | 0.74 |  |
| BW normalized kidney weight | -0.400 | 0.09 |  |
| BW normalized liver weight | 0.003 | 0.99 |  |
| BW normalized spleen weight | -0.356 | **0.004** | UniBa < LUMC |
| Maximal forelimb grip strength | -0.001 | 0.80 |  |
| BW normalized maximal forelimb grip strength | 0.298 | 0.20 |  |

**Supplementary Table 1. Direct site comparisons.**  Outcome measures in which the same SOP was used were directly compared between the two study sites. Differences were very minimal, but significant (indicated in bold) due to the large sample size. BW; body weight.

| **Outcome** | **Site** | **Age**  **(weeks)** | **BL10-WT vs.**  **BL10-*mdx*** | **D2-WT**  **vs. D2-*mdx*** | **BL10-*mdx***  **vs. D2-*mdx*** | **BL10-WT**  **vs. D2-WT** |
| --- | --- | --- | --- | --- | --- | --- |
| **Bodyweight (g)** | LUMC | 2 | 0.31 | 0.28 | 0.26 | 0.48 |
|  | UniBa | 4 | 0.10 | **<0.001** | **<0.001** | **<0.001** |
|  | LUMC | 4 | 0.08 | **0.003** | **<0.001** | **<0.001** |
|  | UniBa | 8 | 0.99 | **<0.001** | **<0.001** | **<0.001** |
|  | LUMC | 8 | **0.042** | **0.001** | **<0.001** | **0.001** |
|  | UniBa | 12 | **<0.001** | **<0.001** | **<0.001** | **<0.001** |
|  | LUMC | 12 | **<0.001** | **<0.001** | **<0.001** | **<0.001** |
|  | UniBa | 16 | 0.09 | **<0.001** | **<0.001** | **<0.001** |
|  | LUMC | 16 | **<0.001** | **<0.001** | **<0.001** | **<0.001** |
|  | UniBa | 18 | 0.06 | **0.005** | **<0.001** | 0.31 |
|  | UniBa | 20 | 0.61 | **0.001** | **<0.001** | **<0.001** |
|  | LUMC | 20 | **0.001** | **<0.001** | **<0.001** | 0.16 |
|  | UniBa | 24 | 0.11 | **<0.001** | **<0.001** | **0.010** |
|  | LUMC | 24 | 0.54 | **<0.001** | **<0.001** | 0.25 |
|  | UniBa | 28 | 0.56 | **<0.001** | **<0.001** | **0.014** |
|  | LUMC | 28 | **0.007** | **<0.001** | **<0.001** | 0.80 |
|  | UniBa | 32 | 0.78 | **<0.001** | **<0.001** | **<0.001** |
|  | LUMC | 32 | 0.99 | **0.001** | **<0.001** | 0.99 |
|  | UniBa | 36 | 0.93 | **<0.001** | **<0.001** | **<0.001** |
|  | LUMC | 36 | 0.99 | **0.022** | **<0.001** | 0.90 |
|  | UniBa | 40 | 0.99 | **<0.001** | **<0.001** | **<0.001** |
|  | LUMC | 40 | 0.89 | **0.007** | **<0.001** | 0.45 |
|  | LUMC | 44 | 0.61 | **0.002** | **<0.001** | 0.57 |
|  | UniBa | 46 | 0.69 | **<0.001** | **<0.001** | **<0.001** |
|  | LUMC | 48 | 0.56 | **0.002** | **<0.001** | 0.77 |
|  | UniBa | 52 | 0.99 | **<0.001** | **<0.001** | **<0.001** |
|  | LUMC | 52 | 0.53 | **0.013** | **<0.001** | 0.92 |

**Supplementary Table 2. Strain comparisons of body weight.** Adjusted *P­*-values of the strain by strain comparisons for each of the study sites. *P*<0.05 is indicated in bold.

| **Outcome** | **Site** | **Age**  **(weeks)** | **BL10-WT vs.**  **BL10-*mdx*** | **D2-WT**  **vs. D2-*mdx*** | **BL10-*mdx***  **vs. D2-*mdx*** | **BL10-WT**  **vs. D2-WT** |
| --- | --- | --- | --- | --- | --- | --- |
| **Lean mass (g)** | LUMC | 4 | 0.99 | **<0.001** | **<0.001** | **<0.001** |
|  | LUMC | 8 | **<0.001** | **<0.001** | **<0.001** | **<0.001** |
|  | LUMC | 12 | **<0.001** | **<0.001** | **<0.001** | **<0.001** |
|  | LUMC | 16 | **<0.001** | **<0.001** | **<0.001** | **<0.001** |
|  | LUMC | 20 | **<0.001** | **<0.001** | **<0.001** | **<0.001** |
|  | LUMC | 24 | **<0.001** | **<0.001** | **<0.001** | **<0.001** |
|  | LUMC | 28 | **<0.001** | **<0.001** | **<0.001** | **<0.001** |
|  | LUMC | 32 | **<0.001** | **<0.001** | **<0.001** | 0.74 |
|  | LUMC | 36 | **<0.001** | **<0.001** | **<0.001** | 0.85 |
|  | LUMC | 40 | **<0.001** | **<0.001** | **<0.001** | 0.94 |
|  | LUMC | 44 | **0.004** | **<0.001** | **<0.001** | 0.97 |
|  | LUMC | 48 | **0.011** | **<0.001** | **<0.001** | 0.85 |
|  | LUMC | 52 | 0.17 | **0.011** | **<0.001** | 0.90 |
| **Lean mass (g/BW)** | LUMC | 4 | **<0.001** | **<0.001** | **<0.001** | **<0.001** |
|  | LUMC | 8 | **0.001** | 0.15 | **<0.001** | **<0.001** |
|  | LUMC | 12 | **<0.001** | 0.71 | **<0.001** | **<0.001** |
|  | LUMC | 16 | **<0.001** | 0.99 | **<0.001** | **0.002** |
|  | LUMC | 20 | **<0.001** | 0.75 | **<0.001** | 0.13 |
|  | LUMC | 24 | **<0.001** | **0.014** | **<0.001** | 0.20 |
|  | LUMC | 28 | **<0.001** | **0.004** | **<0.001** | 0.15 |
|  | LUMC | 32 | **<0.001** | 0.37 | **0.002** | 0.99 |
|  | LUMC | 36 | **<0.001** | 0.45 | **<0.001** | 0.99 |
|  | LUMC | 40 | **<0.001** | 0.98 | 0.06 | 0.26 |
|  | LUMC | 44 | **<0.001** | 0.99 | **<0.001** | 0.73 |
|  | LUMC | 48 | **<0.001** | 0.99 | **0.003** | 0.67 |
|  | LUMC | 52 | **<0.001** | 0.85 | **0.016** | 0.71 |
| **Fat mass (g)** | LUMC | 4 | 0.36 | **<0.001** | **<0.001** | **0.002** |
|  | LUMC | 8 | **0.029** | **0.002** | **<0.001** | **<0.001** |
|  | LUMC | 12 | **<0.001** | 0.99 | **<0.001** | **<0.001** |
|  | LUMC | 16 | **<0.001** | 0.39 | **<0.001** | **0.017** |
|  | LUMC | 20 | **<0.001** | **0.028** | **<0.001** | 0.17 |
|  | LUMC | 24 | **<0.001** | **0.011** | **0.011** | 0.66 |
|  | LUMC | 28 | **<0.001** | **0.002** | 0.09 | 0.79 |
|  | LUMC | 32 | **<0.001** | 0.14 | 0.07 | 0.86 |
|  | LUMC | 36 | **<0.001** | 0.19 | 0.18 | 0.99 |
|  | LUMC | 40 | **<0.001** | 0.65 | 0.45 | 0.41 |
|  | LUMC | 44 | **<0.001** | 0.80 | 0.23 | 0.63 |
|  | LUMC | 48 | **<0.001** | 0.81 | 0.36 | 0.66 |
|  | LUMC | 52 | **<0.001** | 0.44 | 0.35 | 0.65 |
| **Fat mass (g/BW)** | LUMC | 4 | 0.24 | **<0.001** | 0.99 | **<0.001** |
|  | LUMC | 8 | **0.007** | 0.55 | **<0.001** | **<0.001** |
|  | LUMC | 12 | **<0.001** | 0.10 | **<0.001** | **<0.001** |
|  | LUMC | 16 | **<0.001** | 0.99 | **<0.001** | **<0.001** |
|  | LUMC | 20 | **<0.001** | 0.13 | **<0.001** | **0.024** |
|  | LUMC | 24 | **<0.001** | **0.036** | **<0.001** | 0.23 |
|  | LUMC | 28 | **<0.001** | **0.006** | **<0.001** | 0.37 |
|  | LUMC | 32 | **<0.001** | 0.32 | **0.002** | 0.71 |
|  | LUMC | 36 | **<0.001** | 0.41 | **0.017** | 0.99 |
|  | LUMC | 40 | **<0.001** | 0.96 | **0.035** | 0.43 |
|  | LUMC | 44 | **<0.001** | 0.99 | **0.011** | 0.60 |
|  | LUMC | 48 | **<0.001** | 0.99 | **0.036** | 0.62 |
|  | LUMC | 52 | **<0.001** | 0.94 | **0.038** | 0.62 |

**Supplementary Table 3. Strain comparisons of lean and fat mass, absolute or normalized by body weight.** Adjusted *P­*-values of the strain by strain comparisons for each of the study sites. *P*<0.05 is indicated in bold.

| **Outcome** | **Site** | **Age**  **(weeks)** | **BL10-WT vs.**  **BL10-*mdx*** | **D2-WT**  **vs. D2-*mdx*** | **BL10-*mdx***  **vs. D2-*mdx*** | **BL10-WT**  **vs. D2-WT** |
| --- | --- | --- | --- | --- | --- | --- |
| **Quadriceps (mg/g)** | LUMC | 2 | 0.98 | 0.85 | 0.12 | 0.55 |
|  | UniBa | 4 | 0.99 | 0.80 | 0.97 | 0.80 |
|  | LUMC | 4 | 0.37 | **0.023** | **0.031** | 0.94 |
|  | UniBa | 8 | **0.042** | **<0.001** | **<0.001** | **0.009** |
|  | LUMC | 8 | **<0.001** | 0.25 | **<0.001** | 0.43 |
|  | UniBa | 12 | **<0.001** | **<0.001** | **<0.001** | **0.048** |
|  | LUMC | 12 | **<0.001** | **<0.001** | **<0.001** | **0.021** |
|  | UniBa | 28 | **<0.001** | **<0.001** | **<0.001** | **<0.001** |
|  | LUMC | 28 | **<0.001** | **0.003** | **<0.001** | 0.99 |
|  | UniBa | 52 | **<0.001** | **<0.001** | **<0.001** | 0.90 |
|  | LUMC | 52 | **<0.001** | **<0.001** | **<0.001** | 0.13 |
| **Gastrocnemius (mg/g)** | LUMC | 2 | 0.88 | 0.84 | 0.93 | 0.79 |
|  | UniBa | 4 | 0.99 | 0.87 | 0.94 | 0.97 |
|  | LUMC | 4 | **0.012** | **0.008** | 0.87 | 0.77 |
|  | UniBa | 8 | **0.031** | 0.67 | **<0.001** | **<0.001** |
|  | LUMC | 8 | 0.11 | 0.99 | **<0.001** | **0.025** |
|  | UniBa | 12 | **<0.001** | 0.08 | **<0.001** | 0.11 |
|  | LUMC | 12 | **<0.001** | 0.46 | **<0.001** | **0.017** |
|  | UniBa | 28 | 0.43 | **<0.001** | **<0.001** | **<0.001** |
|  | LUMC | 28 | **<0.001** | **<0.001** | **<0.001** | **<0.001** |
|  | UniBa | 52 | 0.51 | **<0.001** | **<0.001** | **0.002** |
|  | LUMC | 52 | 0.99 | **<0.001** | **<0.001** | 0.17 |
| **Tibialis anterior (mg/g)** | LUMC | 2 | 0.99 | 0.95 | 0.15 | 0.48 |
|  | UniBa | 4 | 0.84 | 0.92 | 0.99 | 0.61 |
|  | LUMC | 4 | 0.13 | 0.76 | 0.40 | **0.046** |
|  | UniBa | 8 | **<0.001** | 0.99 | **<0.001** | 0.07 |
|  | LUMC | 8 | **<0.001** | 0.99 | **<0.001** | 0.15 |
|  | UniBa | 12 | **<0.001** | 0.42 | **<0.001** | 0.28 |
|  | LUMC | 12 | **<0.001** | 0.21 | **<0.001** | 0.14 |
|  | UniBa | 28 | **<0.001** | **0.007** | **<0.001** | **0.040** |
|  | LUMC | 28 | **<0.001** | 0.49 | **<0.001** | 0.58 |
|  | UniBa | 52 | **<0.001** | **<0.001** | **<0.001** | 0.99 |
|  | LUMC | 52 | **<0.001** | **0.004** | **<0.001** | 0.75 |
| **Triceps (mg/g)** | LUMC | 2 | 0.97 | 0.99 | 0.99 | 0.68 |
|  | UniBa | 4 | 0.70 | 0.98 | 0.99 | 0.46 |
|  | LUMC | 4 | 0.61 | 0.30 | 0.87 | 0.99 |
|  | UniBa | 8 | **0.026** | 0.94 | **<0.001** | **<0.001** |
|  | LUMC | 8 | **<0.001** | 0.99 | **<0.001** | **0.015** |
|  | UniBa | 12 | **0.002** | **0.027** | **<0.001** | 0.14 |
|  | LUMC | 12 | **<0.001** | **0.006** | **<0.001** | 0.16 |
|  | UniBa | 28 | **0.019** | **0.007** | **<0.001** | 0.09 |
|  | LUMC | 28 | **<0.001** | **0.001** | **<0.001** | 0.99 |
|  | UniBa | 52 | **0.001** | **<0.001** | **<0.001** | 0.99 |
|  | LUMC | 52 | **<0.001** | 0.43 | **<0.001** | 0.53 |
| **EDL (mg/g)** | UniBa | 4 | 0.33 | 0.76 | 0.30 | 0.99 |
|  | UniBa | 8 | 0.21 | 0.75 | **<0.001** | **0.026** |
|  | UniBa | 12 | **0.004** | 0.65 | **<0.001** | 0.99 |
|  | UniBa | 28 | **0.005** | 0.82 | **0.005** | 0.81 |
|  | UniBa | 52 | **0.005** | 0.99 | **0.004** | 0.93 |
| **Heart (mg/g)** | LUMC | 2 | **0.026** | 0.76 | 0.45 | 0.06 |
|  | UniBa | 4 | 0.25 | **<0.001** | **<0.001** | 0.07 |
|  | LUMC | 4 | 0.54 | 0.99 | 0.33 | **0.002** |
|  | UniBa | 8 | 0.48 | **0.006** | **<0.001** | **<0.001** |
|  | LUMC | 8 | 0.86 | 0.99 | 0.20 | 0.88 |
|  | UniBa | 12 | 0.98 | 0.41 | **<0.001** | **0.001** |
|  | LUMC | 12 | 0.57 | 0.41 | 0.10 | 0.97 |
|  | UniBa | 28 | 0.98 | 0.23 | **0.005** | **0.003** |
|  | LUMC | 28 | 0.85 | 0.28 | **0.027** | 0.99 |
|  | UniBa | 52 | **0.006** | **0.004** | **<0.001** | **<0.001** |
|  | LUMC | 52 | 0.46 | 0.14 | **<0.001** | 0.70 |

**Supplementary Table 4. Strain comparisons of muscle weights normalized by body weight.** Adjusted *P­*-values of the strain by strain comparisons for each of the study sites. *P*<0.05 is indicated in bold.

| **Outcome** | **Site** | **Age**  **(weeks)** | **BL10-WT vs.**  **BL10-*mdx*** | **D2-WT**  **vs. D2-*mdx*** | **BL10-*mdx***  **vs. D2-*mdx*** | **BL10-WT**  **vs. D2-WT** |
| --- | --- | --- | --- | --- | --- | --- |
| **Kidney (mg/g)** | LUMC | 2 | 0.99 | 0.22 | **<0.001** | **<0.001** |
|  | UniBa | 4 | 0.09 | **0.029** | **0.003** | **0.021** |
|  | LUMC | 4 | 0.99 | **0.035** | 0.61 | 0.52 |
|  | UniBa | 8 | 0.88 | **0.003** | **<0.001** | **0.040** |
|  | LUMC | 8 | 0.81 | 0.79 | **0.021** | 0.30 |
|  | UniBa | 12 | 0.99 | 0.62 | **<0.001** | **<0.001** |
|  | LUMC | 12 | 0.14 | **0.033** | **<0.001** | **<0.001** |
|  | UniBa | 28 | 0.99 | 0.052 | **<0.001** | **<0.001** |
|  | LUMC | 28 | 0.53 | **0.014** | **<0.001** | **<0.001** |
|  | UniBa | 52 | **<0.001** | 0.15 | **<0.001** | **<0.001** |
|  | LUMC | 52 | 0.46 | **<0.001** | **<0.001** | 0.99 |
| **Liver (mg/g)** | LUMC | 2 | 0.98 | 0.77 | 0.99 | 0.65 |
|  | UniBa | 4 | 0.43 | 0.42 | 0.73 | 0.09 |
|  | LUMC | 4 | 0.99 | 0.06 | 0.92 | 0.33 |
|  | UniBa | 8 | 0.88 | **<0.001** | 0.91 | **0.004** |
|  | LUMC | 8 | **<0.001** | 0.56 | 0.11 | 0.43 |
|  | UniBa | 12 | **0.005** | 0.50 | 0.96 | 0.09 |
|  | LUMC | 12 | **0.018** | **<0.001** | **0.025** | 0.37 |
|  | UniBa | 28 | 0.40 | 0.31 | 0.59 | 0.08 |
|  | LUMC | 28 | 0.79 | **<0.001** | 0.11 | 0.99 |
|  | UniBa | 52 | **<0.001** | 0.99 | 0.98 | **<0.001** |
|  | LUMC | 52 | **0.038** | **0.039** | **0.001** | **<0.001** |
| **Spleen (mg/g)** | LUMC | 2 | 0.33 | 0.95 | **<0.001** | **0.040** |
|  | UniBa | 4 | 0.10 | **0.003** | **0.033** | **<0.001** |
|  | LUMC | 4 | 0.92 | 0.65 | 0.86 | 0.72 |
|  | UniBa | 8 | 0.97 | 0.63 | 0.91 | 0.99 |
|  | LUMC | 8 | **<0.001** | 0.06 | 0.99 | 0.55 |
|  | UniBa | 12 | 0.23 | 0.94 | 0.99 | 0.23 |
|  | LUMC | 12 | **0.001** | 0.50 | 0.95 | 0.60 |
|  | UniBa | 28 | 0.99 | **0.037** | **0.020** | 0.98 |
|  | LUMC | 28 | 0.99 | **0.009** | 0.17 | 0.98 |
|  | UniBa | 52 | 0.42 | 0.99 | **0.008** | 0.71 |
|  | LUMC | 52 | 0.09 | 0.88 | 0.34 | **0.033** |
| **Tibia length (mm)** | LUMC | 2 | 0.38 | 0.47 | 0.95 | 0.09 |
|  | LUMC | 4 | 0.33 | 0.15 | 0.99 | 0.80 |
|  | UniBa | 8 | 0.12 | 0.07 | 0.78 | 0.65 |
|  | LUMC | 8 | 0.99 | **<0.001** | **0.003** | 0.89 |
|  | UniBa | 12 | 0.85 | 0.94 | 0.99 | 0.99 |
|  | LUMC | 12 | 0.99 | 0.85 | 0.92 | 0.99 |
|  | UniBa | 28 | 0.89 | 0.82 | 0.97 | 0.96 |
|  | LUMC | 28 | **0.046** | 0.61 | 0.76 | 0.99 |
|  | UniBa | 52 | 0.99 | 0.62 | 0.90 | 0.99 |
|  | LUMC | 52 | 0.09 | 0.96 | 0.99 | **0.011** |
| **Femur length (mm)** | UniBa | 4 | 0.99 | **<0.001** | 0.16 | 0.09 |
|  | UniBa | 8 | 0.56 | 0.83 | 0.69 | 0.26 |
|  | UniBa | 12 | 0.99 | 0.99 | 0.99 | 0.99 |
|  | UniBa | 28 | 0.29 | 0.29 | 0.29 | 0.34 |
|  | UniBa | 52 | 0.99 | 0.99 | 0.92 | 0.99 |

**Supplementary Table 5. Strain comparisons of organ weights normalized by body weight and bone length.** Adjusted *P­*-values of the strain by strain comparisons for each of the study sites. *P*<0.05 is indicated in bold.

| **Outcome** | **Site** | **Age**  **(weeks)** | **BL10-WT vs.**  **BL10-*mdx*** | **D2-WT**  **vs. D2-*mdx*** | **BL10-*mdx***  **vs. D2-*mdx*** | **BL10-WT**  **vs. D2-WT** |
| --- | --- | --- | --- | --- | --- | --- |
| **Absolute forelimb grip strength (KgF)** | UniBa | 4 | **<0.001** | **<0.001** | **0.008** | **<0.001** |
|  | LUMC | 4 | **0.006** | **<0.001** | 0.06 | 0.98 |
|  | UniBa | 8 | **<0.001** | **<0.001** | **<0.001** | **0.042** |
|  | LUMC | 8 | **<0.001** | **<0.001** | 0.53 | 0.99 |
|  | UniBa | 12 | **<0.001** | **<0.001** | **<0.001** | **<0.001** |
|  | LUMC | 12 | **0.002** | **<0.001** | 0.85 | 0.17 |
|  | UniBa | 16 | **<0.001** | **<0.001** | **<0.001** | **0.030** |
|  | LUMC | 16 | 0.98 | **<0.001** | 0.74 | 0.05 |
|  | UniBa | 18 | 0.43 | **<0.001** | 0.37 | 0.57 |
|  | UniBa | 20 | **0.044** | **<0.001** | **<0.001** | 0.19 |
|  | LUMC | 20 | 0.25 | **<0.001** | 0.85 | 0.06 |
|  | UniBa | 24 | **<0.001** | **<0.001** | **<0.001** | 0.77 |
|  | LUMC | 24 | 0.11 | **<0.001** | 0.93 | **0.026** |
|  | UniBa | 28 | **<0.001** | **<0.001** | **<0.001** | **0.007** |
|  | LUMC | 28 | **0.022** | 0.48 | **0.042** | 0.99 |
|  | UniBa | 32 | **<0.001** | **<0.001** | **<0.001** | **0.006** |
|  | LUMC | 32 | **0.007** | 0.06 | **0.048** | 0.97 |
|  | UniBa | 36 | **<0.001** | **<0.001** | **<0.001** | **0.018** |
|  | LUMC | 36 | **0.039** | **<0.001** | 0.93 | 0.85 |
|  | UniBa | 40 | **<0.001** | **<0.001** | **<0.001** | **<0.001** |
|  | LUMC | 40 | 0.99 | **0.011** | 0.96 | 0.20 |
|  | LUMC | 44 | **0.013** | **<0.001** | 0.27 | 0.20 |
|  | UniBa | 46 | **<0.001** | **<0.001** | **0.008** | **0.003** |
|  | LUMC | 48 | 0.06 | **0.002** | 0.96 | 0.83 |
|  | UniBa | 52 | **0.014** | **<0.001** | **<0.001** | 0.99 |
|  | LUMC | 52 | **0.007** | **<0.001** | 0.87 | 0.44 |
| **BW normalized forelimb grip strength (KgF/Kg)** | UniBa | 4 | **<0.001** | 0.98 | **<0.001** | **0.046** |
|  | LUMC | 4 | **0.003** | **0.036** | **0.026** | 0.69 |
|  | UniBa | 8 | **<0.001** | **0.009** | **0.018** | **0.002** |
|  | LUMC | 8 | **<0.001** | **0.007** | **<0.001** | 0.47 |
|  | UniBa | 12 | **<0.001** | **<0.001** | 0.06 | 0.99 |
|  | LUMC | 12 | **<0.001** | **<0.001** | **<0.001** | **0.011** |
|  | UniBa | 16 | **<0.001** | **<0.001** | 0.16 | 0.33 |
|  | LUMC | 16 | 0.34 | 0.55 | **0.016** | 0.15 |
|  | UniBa | 18 | 0.11 | **0.002** | **0.001** | 0.08 |
|  | UniBa | 20 | **0.013** | 0.05 | 0.88 | 0.21 |
|  | LUMC | 20 | **0.019** | 0.07 | **<0.001** | **0.002** |
|  | UniBa | 24 | **<0.001** | **0.004** | **0.006** | 0.24 |
|  | LUMC | 24 | **0.024** | **0.035** | **<0.001** | **0.002** |
|  | UniBa | 28 | **<0.001** | **<0.001** | 0.76 | 0.99 |
|  | LUMC | 28 | **<0.001** | 0.26 | **<0.001** | 0.88 |
|  | UniBa | 32 | **<0.001** | **0.002** | 0.07 | 0.94 |
|  | LUMC | 32 | **0.004** | 0.99 | **<0.001** | 0.86 |
|  | UniBa | 36 | **0.002** | **<0.001** | 0.99 | 0.94 |
|  | LUMC | 36 | 0.15 | 0.10 | 0.83 | 0.27 |
|  | UniBa | 40 | **<0.001** | **<0.001** | 0.51 | 0.43 |
|  | LUMC | 40 | 0.99 | 0.68 | 0.17 | 0.08 |
|  | UniBa | 46 | **0.001** | **<0.001** | 0.29 | 0.27 |
|  | LUMC | 44 | **0.012** | 0.16 | **0.003** | **0.040** |
|  | LUMC | 48 | 0.09 | 0.18 | **0.026** | 0.40 |
|  | UniBa | 52 | 0.11 | **0.002** | 0.56 | **0.004** |
|  | LUMC | 52 | **0.039** | **0.031** | 0.97 | 0.12 |

**Supplementary Table 6. Strain comparisons of forelimb grip strength.** Adjusted *P­*-values of the strain by strain comparisons for each of the study sites. *P*<0.05 is indicated in bold.

| **Outcome** | **Site** | **Age**  **(weeks)** | **BL10-WT vs.**  **BL10-*mdx*** | **D2-WT**  **vs. D2-*mdx*** | **BL10-*mdx***  **vs. D2-*mdx*** | **BL10-WT**  **vs. D2-WT** |
| --- | --- | --- | --- | --- | --- | --- |
| **Maximum hang time (s)** | LUMC | 4 | **<0.001** | **<0.001** | 0.83 | 0.30 |
|  | LUMC | 8 | **<0.001** | **<0.001** | 0.80 | 0.78 |
|  | LUMC | 12 | **<0.001** | **<0.001** | **<0.001** | 0.93 |
|  | LUMC | 16 | **<0.001** | **<0.001** | **0.004** | 0.99 |
|  | LUMC | 20 | **<0.001** | **<0.001** | 0.05 | 0.22 |
|  | LUMC | 24 | **<0.001** | **<0.001** | 0.18 | 0.39 |
|  | LUMC | 28 | **<0.001** | **0.007** | 0.47 | 0.30 |
|  | LUMC | 32 | **<0.001** | 0.07 | 0.50 | **0.027** |
|  | LUMC | 36 | **<0.001** | **0.042** | 0.76 | **0.014** |
|  | LUMC | 40 | **<0.001** | **0.002** | 0.59 | **0.002** |
|  | LUMC | 44 | **<0.001** | **0.006** | 0.79 | **<0.001** |
|  | LUMC | 48 | **<0.001** | **0.001** | 0.54 | **0.005** |
|  | LUMC | 52 | **<0.001** | **0.014** | 0.77 | **0.011** |
| **Total distance moved (cm)** | LUMC | 4 | 0.99 | **<0.001** | **<0.001** | 0.12 |
|  | LUMC | 8 | 0.17 | **<0.001** | **<0.001** | 0.22 |
|  | LUMC | 12 | **0.004** | **<0.001** | 0.29 | 0.87 |
|  | LUMC | 16 | 0.20 | **0.001** | 0.75 | **0.017** |
|  | LUMC | 20 | 0.10 | **<0.001** | 0.38 | **0.023** |
|  | LUMC | 24 | 0.12 | **<0.001** | **0.005** | **<0.001** |
|  | LUMC | 28 | **0.014** | **0.028** | **0.009** | **0.004** |
|  | LUMC | 32 | 0.35 | **0.006** | 0.08 | **<0.001** |
|  | LUMC | 36 | 0.05 | **0.029** | **0.016** | **<0.001** |
|  | LUMC | 40 | 0.06 | 0.13 | **0.003** | **<0.001** |
|  | LUMC | 44 | **0.002** | 0.67 | 0.07 | **0.005** |
|  | LUMC | 48 | 0.66 | 0.17 | **0.044** | **<0.001** |
|  | LUMC | 52 | 0.38 | 0.89 | **0.012** | **0.017** |
| **Total time moving (s)** | LUMC | 4 | 0.89 | **<0.001** | **<0.001** | **0.003** |
|  | LUMC | 8 | 0.82 | **<0.001** | **<0.001** | **0.044** |
|  | LUMC | 12 | **0.006** | **<0.001** | **0.033** | 0.94 |
|  | LUMC | 16 | 0.16 | **0.001** | 0.99 | 0.17 |
|  | LUMC | 20 | 0.08 | **<0.001** | 0.73 | 0.14 |
|  | LUMC | 24 | 0.11 | **<0.001** | **0.006** | **<0.001** |
|  | LUMC | 28 | **0.013** | **0.022** | **0.010** | **0.015** |
|  | LUMC | 32 | 0.48 | **0.007** | 0.11 | **<0.001** |
|  | LUMC | 36 | 0.08 | 0.12 | **0.048** | 0.13 |
|  | LUMC | 40 | **0.048** | **<0.001** | **0.037** | 0.26 |
|  | LUMC | 44 | **0.002** | 0.09 | **0.015** | **0.002** |
|  | LUMC | 48 | 0.83 | 0.38 | **0.026** | **<0.001** |
|  | LUMC | 52 | 0.43 | 0.93 | **0.005** | **0.007** |
| **Mean velocity (cm/s)** | LUMC | 4 | 0.99 | **<0.001** | **<0.001** | 0.09 |
|  | LUMC | 8 | 0.10 | **<0.001** | **<0.001** | 0.16 |
|  | LUMC | 12 | **0.004** | **<0.001** | 0.29 | 0.78 |
|  | LUMC | 16 | 0.16 | **0.001** | 0.71 | **0.013** |
|  | LUMC | 20 | 0.10 | **<0.001** | 0.39 | **0.019** |
|  | LUMC | 24 | 0.11 | **<0.001** | **0.005** | **<0.001** |
|  | LUMC | 28 | **0.015** | **0.026** | **0.010** | **0.004** |
|  | LUMC | 32 | 0.35 | **0.006** | 0.08 | **<0.001** |
|  | LUMC | 36 | **0.045** | **0.028** | **0.015** | **<0.001** |
|  | LUMC | 40 | **0.060** | **0.018** | **0.004** | **<0.001** |
|  | LUMC | 44 | **0.002** | 0.74 | 0.08 | **0.006** |
|  | LUMC | 48 | 0.69 | 0.15 | **0.045** | **<0.001** |
|  | LUMC | 52 | 0.38 | 0.85 | **0.011** | **0.015** |
| **Time inner zone (s)** | LUMC | 4 | 0.94 | 0.41 | 0.92 | **0.049** |
|  | LUMC | 8 | 0.06 | 0.99 | 0.95 | 0.28 |
|  | LUMC | 12 | **0.038** | 0.99 | 0.67 | 0.97 |
|  | LUMC | 16 | 0.15 | 0.43 | 0.10 | **0.021** |
|  | LUMC | 20 | 0.36 | 0.96 | 0.45 | 0.22 |
|  | LUMC | 24 | 0.29 | 0.23 | 0.07 | **0.014** |
|  | LUMC | 28 | 0.53 | 0.88 | 0.36 | 0.94 |
|  | LUMC | 32 | 0.96 | 0.14 | 0.28 | **0.024** |
|  | LUMC | 36 | 0.88 | 0.95 | 0.06 | **0.001** |
|  | LUMC | 40 | 0.98 | 0.98 | 0.19 | 0.09 |
|  | LUMC | 44 | 0.94 | 0.98 | 0.27 | **0.046** |
|  | LUMC | 48 | 0.54 | 0.91 | 0.018 | **0.008** |
|  | LUMC | 52 | 0.91 | 0.59 | 0.025 | **0.006** |

**Supplementary Table 7. Strain comparisons of two limb hanging test and open field test.** Adjusted *P­*-values of the strain by strain comparisons for each of the study sites. *P*<0.05 is indicated in bold.

| **Outcome** | **Site** | **Age**  **(weeks)** | **BL10-WT vs.**  **BL10-*mdx*** | **D2-WT**  **vs. D2-*mdx*** | **BL10-*mdx***  **vs. D2-*mdx*** | **BL10-WT**  **vs. D2-WT** |
| --- | --- | --- | --- | --- | --- | --- |
| **Respiratory rate / s** | LUMC | 4 | 0.12 | **<0.001** | 0.99 | 0.11 |
|  | LUMC | 8 | **<0.001** | **<0.001** | **0.002** | **0.021** |
|  | LUMC | 12 | **<0.001** | **<0.001** | **<0.001** | **<0.001** |
|  | LUMC | 16 | **<0.001** | 0.12 | **<0.001** | **<0.001** |
|  | LUMC | 20 | **0.004** | 0.08 | **<0.001** | **<0.001** |
|  | LUMC | 24 | **0.003** | 0.49 | **<0.001** | **<0.001** |
|  | LUMC | 28 | **0.010** | 0.12 | **<0.001** | **<0.001** |
|  | LUMC | 32 | **0.022** | 0.94 | **<0.001** | **<0.001** |
|  | LUMC | 36 | **0.029** | 0.81 | **<0.001** | **0.004** |
|  | LUMC | 40 | 0.78 | 0.92 | **<0.001** | **<0.001** |
|  | LUMC | 44 | 0.33 | 0.69 | **<0.001** | **<0.001** |
|  | LUMC | 48 | 0.71 | 0.99 | **<0.001** | **0.011** |
|  | LUMC | 52 | 0.99 | 0.99 | **<0.001** | **<0.001** |
| **Respiratory amplitude (mV)** | LUMC | 4 | 0.03 | **<0.001** | **<0.001** | **<0.001** |
|  | LUMC | 8 | 0.11 | **<0.001** | **<0.001** | **<0.001** |
|  | LUMC | 12 | **0.041** | **<0.001** | **<0.001** | **0.006** |
|  | LUMC | 16 | 0.62 | **<0.001** | **<0.001** | 0.60 |
|  | LUMC | 20 | **0.034** | **0.009** | 0.92 | 0.91 |
|  | LUMC | 24 | **0.003** | **<0.001** | 0.84 | 0.42 |
|  | LUMC | 28 | **0.002** | **0.016** | 0.68 | 0.97 |
|  | LUMC | 32 | **0.024** | 0.17 | 0.92 | 0.75 |
|  | LUMC | 36 | 0.99 | **0.021** | 0.95 | **0.002** |
|  | LUMC | 40 | 0.06 | 0.99 | 0.76 | 0.75 |
|  | LUMC | 44 | **0.021** | **0.025** | 0.08 | 0.73 |
|  | LUMC | 48 | **<0.001** | **0.003** | 0.99 | 0.73 |
|  | LUMC | 52 | **<0.001** | 0.23 | 0.09 | 0.93 |
| **Respiratory amplitude (mV/g)** | LUMC | 4 | 0.24 | 0.99 | 0.97 | 0.07 |
|  | LUMC | 8 | **0.013** | **0.044** | 0.75 | **0.039** |
|  | LUMC | 12 | **<0.001** | **0.001** | 0.99 | 0.67 |
|  | LUMC | 16 | **0.009** | **0.003** | 0.79 | 0.99 |
|  | LUMC | 20 | **0.002** | 0.99 | **<0.001** | 0.99 |
|  | LUMC | 24 | **<0.001** | 0.83 | **0.003** | 0.99 |
|  | LUMC | 28 | **<0.001** | 0.36 | **<0.001** | 0.98 |
|  | LUMC | 32 | **0.012** | 0.58 | **0.018** | 0.85 |
|  | LUMC | 36 | **0.027** | 0.99 | **<0.001** | 0.48 |
|  | LUMC | 40 | 0.06 | 0.25 | **0.003** | 0.99 |
|  | LUMC | 44 | **0.045** | 0.85 | **<0.001** | 0.06 |
|  | LUMC | 48 | **0.001** | 0.99 | **0.027** | 0.99 |
|  | LUMC | 52 | **0.003** | 0.53 | **<0.001** | 0.86 |

**Supplementary Table 8. Strain comparisons of respiratory functionality.** Adjusted *P­*-values of the strain by strain comparisons for each of the study sites. *P*<0.05 is indicated in bold.

| **Outcome** | **Site** | **Frequency**  **(Hz)** | **BL10-WT vs.**  **BL10-*mdx*** | **D2-WT**  **vs. D2-*mdx*** | **BL10-*mdx***  **vs. D2-*mdx*** | **BL10-WT**  **vs. D2-WT** |
| --- | --- | --- | --- | --- | --- | --- |
| **Plantar flexor torque (N*mm/kg)**  **8 weeks of age** | UniBa | 1 | **<0.001** | **<0.001** | **0.034** | 0.06 |
|  | UniBa | 10 | **<0.001** | **<0.001** | **0.016** | 0.07 |
|  | UniBa | 30 | **<0.001** | **0.002** | 0.74 | 0.12 |
|  | UniBa | 50 | **<0.001** | **<0.001** | **0.002** | **0.014** |
|  | UniBa | 80 | **<0.001** | **<0.001** | **<0.001** | **0.007** |
|  | UniBa | 100 | **<0.001** | **<0.001** | **<0.001** | **0.008** |
|  | UniBa | 120 | **<0.001** | **<0.001** | **<0.001** | **0.009** |
|  | UniBa | 150 | **<0.001** | **<0.001** | **<0.001** | **0.008** |
|  | UniBa | 180 | **<0.001** | **<0.001** | **<0.001** | **0.005** |
|  | UniBa | 200 | **<0.001** | **<0.001** | **<0.001** | **0.004** |
| **Plantar flexor torque (N*mm/kg)**  **12 weeks of age** | UniBa | 1 | **<0.001** | **<0.001** | **<0.001** | **0.018** |
|  | UniBa | 10 | **<0.001** | **<0.001** | **<0.001** | **0.008** |
|  | UniBa | 30 | **<0.001** | **<0.001** | **0.048** | 0.47 |
|  | UniBa | 50 | **<0.001** | **<0.001** | **<0.001** | **<0.001** |
|  | UniBa | 80 | **<0.001** | **<0.001** | **<0.001** | **<0.001** |
|  | UniBa | 100 | **<0.001** | **<0.001** | **<0.001** | **<0.001** |
|  | UniBa | 120 | **<0.001** | **<0.001** | **<0.001** | **<0.001** |
|  | UniBa | 150 | **<0.001** | **<0.001** | **<0.001** | **<0.001** |
|  | UniBa | 180 | **<0.001** | **<0.001** | **<0.001** | **<0.001** |
|  | UniBa | 200 | **<0.001** | **<0.001** | **<0.001** | **<0.001** |
| **Plantar flexor torque (N*mm/kg)**  **28 weeks of age** | UniBa | 1 | **<0.001** | **<0.001** | **0.007** | 0.99 |
|  | UniBa | 10 | **<0.001** | **<0.001** | **0.002** | 0.99 |
|  | UniBa | 30 | **<0.001** | **<0.001** | 0.89 | 0.98 |
|  | UniBa | 50 | **<0.001** | **<0.001** | **<0.001** | 0.31 |
|  | UniBa | 80 | **<0.001** | **<0.001** | **<0.001** | **0.002** |
|  | UniBa | 100 | **<0.001** | **<0.001** | **<0.001** | **0.001** |
|  | UniBa | 120 | **<0.001** | **<0.001** | **<0.001** | **<0.001** |
|  | UniBa | 150 | **<0.001** | **<0.001** | **<0.001** | **<0.001** |
|  | UniBa | 180 | **<0.001** | **<0.001** | **<0.001** | **<0.001** |
|  | UniBa | 200 | **<0.001** | **<0.001** | **<0.001** | **<0.001** |
| **Plantar flexor torque (N*mm/kg)**  **52 weeks of age** | UniBa | 1 | **<0.001** | **<0.001** | **<0.001** | **<0.001** |
|  | UniBa | 10 | **<0.001** | **<0.001** | **<0.001** | **0.002** |
|  | UniBa | 30 | **<0.001** | **<0.001** | **<0.001** | 0.94 |
|  | UniBa | 50 | **<0.001** | **<0.001** | **<0.001** | 0.18 |
|  | UniBa | 80 | **<0.001** | **<0.001** | **<0.001** | **0.001** |
|  | UniBa | 100 | **<0.001** | **<0.001** | **<0.001** | **<0.001** |
|  | UniBa | 120 | **<0.001** | **<0.001** | **<0.001** | **<0.001** |
|  | UniBa | 150 | **<0.001** | **<0.001** | **<0.001** | **<0.001** |
|  | UniBa | 180 | **<0.001** | **<0.001** | **<0.001** | **<0.001** |
|  | UniBa | 200 | **<0.001** | **<0.001** | **<0.001** | **<0.001** |

**Supplementary Table 9. Strain comparisons of plantar flexor torque.** Adjusted *P­*-values of the strain by strain comparisons for each of the study sites. *P*<0.05 is indicated in bold.

| **Outcome** | **Site** | **Age**  **(weeks)** | **BL10-WT vs.**  **BL10-*mdx*** | **D2-WT**  **vs. D2-*mdx*** | **BL10-*mdx***  **vs. D2-*mdx*** | **BL10-WT**  **vs. D2-WT** |
| --- | --- | --- | --- | --- | --- | --- |
| **Hindlimb volume (mm^3^)** | UniBa | 8 | 0.19 | **<0.001** | **<0.001** | **0.014** |
|  | UniBa | 12 | 0.55 | **<0.001** | **<0.001** | 0.19 |
|  | UniBa | 28 | **0.024** | **<0.001** | **<0.001** | 0.60 |
|  | UniBa | 52 | 0.99 | **<0.001** | **<0.001** | **0.049** |
| **Gastrocnemius echodensity** | UniBa | 8 | 0.20 | **0.001** | **0.010** | 0.98 |
|  | UniBa | 12 | 0.10 | **<0.001** | **0.002** | **0.002** |
|  | UniBa | 28 | 0.76 | **<0.001** | **0.001** | 0.99 |
|  | UniBa | 52 | 0.94 | **<0.001** | **<0.001** | 0.66 |
| **Diaphragm amplitude (mm)** | UniBa | 8 | 0.05 | **<0.001** | 0.86 | 0.99 |
|  | UniBa | 12 | **<0.001** | **0.006** | **<0.001** | **<0.001** |
|  | UniBa | 28 | **0.003** | **<0.001** | **0.002** | **<0.001** |
|  | UniBa | 52 | 0.09 | **0.041** | **<0.001** | **0.006** |
| **Diaphragm echodensity** | UniBa | 8 | 0.85 | **0.005** | 0.96 | 0.54 |
|  | UniBa | 12 | **0.022** | **0.027** | 0.99 | 0.99 |
|  | UniBa | 28 | **0.011** | 0.99 | 0.24 | 0.78 |
|  | UniBa | 52 | **0.002** | 0.99 | 0.41 | 0.89 |

**Supplementary Table 10. Strain comparisons of muscle echo sonography.** Adjusted *P­*-values of the strain by strain comparisons for each of the study sites. *P*<0.05 is indicated in bold.

| **Outcome** | **Site** | **Age**  **(weeks)** | **BL10-WT vs.**  **BL10-*mdx*** | **D2-WT**  **vs. D2-*mdx*** | **BL10-*mdx***  **vs. D2-*mdx*** | **BL10-WT**  **vs. D2-WT** |
| --- | --- | --- | --- | --- | --- | --- |
| **Stroke volume (µl)**  **Echo** | UniBa | 8 | 0.87 | 0.96 | 0.74 | 0.99 |
|  | UniBa | 12 | 0.95 | 0.74 | 0.80 | 0.95 |
|  | UniBa | 28 | 0.46 | **0.002** | 0.99 | 0.43 |
|  | UniBa | 52 | 0.55 | 0.93 | 0.91 | 0.87 |
| **Stroke volume (µl)**  **MRI** | LUMC | 8 | 0.99 | **0.002** | **<0.001** | 0.99 |
|  | LUMC | 12 | 0.97 | 0.16 | 0.73 | 0.81 |
|  | LUMC | 28 | 0.99 | **0.029** | 0.10 | 0.91 |
|  | LUMC | 52 | **0.003** | **<0.001** | 0.08 | 0.05 |
| **Cardiac output (mL/min)**  **Echo** | UniBa | 8 | 0.99 | 0.62 | 0.47 | 0.97 |
|  | UniBa | 12 | 0.75 | 0.47 | 0.73 | 0.83 |
|  | UniBa | 28 | 0.52 | **0.031** | 0.67 | 0.93 |
|  | UniBa | 52 | 0.62 | 0.60 | 0.27 | 0.85 |
| **Cardiac output (mL/min)**  **MRI** | LUMC | 8 | 0.99 | 0.21 | 0.06 | 0.20 |
|  | LUMC | 12 | 0.99 | 0.48 | 0.20 | 0.29 |
|  | LUMC | 28 | 0.98 | **0.012** | 0.09 | 0.99 |
|  | LUMC | 52 | **0.005** | **0.003** | 0.63 | 0.89 |
| **Ejection fraction (%)**  **Echo** | UniBa | 8 | 0.74 | 0.99 | 0.99 | 0.58 |
|  | UniBa | 12 | 0.99 | 0.20 | 0.17 | 0.99 |
|  | UniBa | 28 | 0.88 | **0.013** | **0.010** | 0.99 |
|  | UniBa | 52 | **<0.001** | **0.028** | 0.95 | **0.028** |
| **Ejection fraction (%)**  **MRI** | LUMC | 8 | 0.93 | **0.013** | **0.002** | 0.99 |
|  | LUMC | 12 | 0.88 | 0.35 | 0.64 | 0.99 |
|  | LUMC | 28 | 0.06 | **0.008** | 0.52 | 0.26 |
|  | LUMC | 52 | **<0.001** | **<0.001** | 0.44 | 0.75 |
| **Shortening fraction (%)**  **Echo** | UniBa | 8 | 0.73 | 0.99 | 0.99 | 0.50 |
|  | UniBa | 12 | 0.99 | 0.15 | 0.13 | 0.99 |
|  | UniBa | 28 | 0.84 | **0.014** | **0.018** | 0.99 |
|  | UniBa | 52 | **<0.001** | **0.021** | 0.90 | **0.026** |

**Supplementary Table 11. Strain comparisons of cardiac outcome measures.** Adjusted *P­*-values of the strain by strain comparisons for each of the study sites. *P*<0.05 is indicated in bold. MRI; magnetic resonance imaging.

| **Outcome** | **Site** | **Age**  **(weeks)** | **BL10-WT vs.**  **BL10-*mdx*** | **D2-WT**  **vs. D2-*mdx*** | **BL10-*mdx***  **vs. D2-*mdx*** | **BL10-WT**  **vs. D2-WT** |
| --- | --- | --- | --- | --- | --- | --- |
| **EDL twitch TTP (ms)** | UniBa | 8 | 0.46 | 0.51 | 0.84 | 0.19 |
|  | UniBa | 12 | 0.86 | 0.99 | 0.41 | 0.37 |
|  | UniBa | 28 | 0.74 | 0.80 | 0.68 | 0.58 |
|  | UniBa | 52 | 0.88 | 0.96 | 0.27 | 0.99 |
| **EDL twitch HRT (ms)** | UniBa | 8 | 0.21 | 0.98 | 0.78 | 0.55 |
|  | UniBa | 12 | 0.61 | 0.56 | 0.67 | 0.64 |
|  | UniBa | 28 | 0.38 | 0.97 | 0.38 | 0.94 |
|  | UniBa | 52 | 0.98 | 0.83 | 0.94 | 0.99 |
| **EDL twitch specific force (kN/m^2^)** | UniBa | 8 | **0.001** | **0.025** | 0.80 | 0.99 |
|  | UniBa | 12 | **<0.001** | **0.032** | 0.19 | 0.022 |
|  | UniBa | 28 | **<0.001** | **0.009** | 0.99 | 0.25 |
|  | UniBa | 52 | **<0.001** | 0.08 | 0.97 | 0.28 |
| **EDL tetanic specific force (kN/m^2^)** | UniBa | 8 | **<0.001** | **0.009** | 0.92 | 0.94 |
|  | UniBa | 12 | **<0.001** | **0.001** | 0.36 | 0.40 |
|  | UniBa | 28 | **<0.001** | **0.006** | 0.99 | 0.37 |
|  | UniBa | 52 | **<0.001** | **0.012** | 0.49 | 0.42 |
| **EDL twitch maximal force (mN)** | UniBa | 8 | 0.56 | **<0.001** | **<0.001** | **0.008** |
|  | UniBa | 12 | 0.11 | **0.003** | **<0.001** | **0.001** |
|  | UniBa | 28 | **0.049** | **0.003** | **0.015** | 0.07 |
|  | UniBa | 52 | **0.007** | **0.004** | **<0.001** | 0.05 |
| **EDL tetanic maximal force (mN)** | UniBa | 8 | **0.020** | **<0.001** | **<0.001** | 0.64 |
|  | UniBa | 12 | 0.99 | **<0.001** | **<0.001** | 0.85 |
|  | UniBa | 28 | 0.12 | **0.001** | **0.004** | **0.036** |
|  | UniBa | 52 | 0.67 | **<0.001** | **<0.001** | **0.003** |
| **EDL force drop at 5^th^ pulse (%)** | UniBa | 8 | 0.30 | 0.99 | 0.45 | 0.99 |
|  | UniBa | 12 | 0.71 | 0.72 | 0.80 | **0.006** |
|  | UniBa | 28 | **0.019** | 0.35 | 0.20 | 0.74 |
|  | UniBa | 52 | **<0.001** | 0.74 | 0.10 | 0.73 |
| **EDL force drop at 10^th^ pulse (%)** | UniBa | 8 | **0.021** | 0.63 | 0.11 | 0.9 |
|  | UniBa | 12 | 0.95 | 0.23 | 0.64 | **0.049** |
|  | UniBa | 28 | 0.11 | **<0.001** | 0.99 | 0.62 |
|  | UniBa | 52 | **<0.001** | **0.026** | 0.43 | 0.67 |
| **EDL force recovery at 5 min (%)** | UniBa | 8 | 0.11 | **0.037** | 0.58 | **0.010** |
|  | UniBa | 12 | **<0.001** | 0.070 | **0.032** | 0.60 |
|  | UniBa | 28 | **0.028** | **<0.001** | 0.96 | 0.65 |
|  | UniBa | 52 | **<0.001** | **0.001** | 0.06 | 0.80 |
| **EDL force recovery at 15 min (%)** | UniBa | 8 | **0.015** | **0.004** | 0.69 | 0.22 |
|  | UniBa | 12 | 0.42 | 0.06 | 0.58 | 0.08 |
|  | UniBa | 28 | **0.006** | **0.001** | 0.99 | 0.52 |
|  | UniBa | 52 | **<0.001** | **0.026** | 0.12 | 0.75 |

**Supplementary Table 12. Strain comparisons of *ex vivo* physiology of the EDL.** Adjusted *P­*-values of the strain by strain comparisons for each of the study sites. *P*<0.05 is indicated in bold.

| **Outcome** | **Site** | **Age**  **(weeks)** | **BL10-WT vs.**  **BL10-*mdx*** | **D2-WT**  **vs. D2-*mdx*** | **BL10-*mdx***  **vs. D2-*mdx*** | **BL10-WT**  **vs. D2-WT** |
| --- | --- | --- | --- | --- | --- | --- |
| **Diaphragm twitch TTP (ms)** | UniBa | 8 | 0.19 | 0.68 | 0.15 | 0.74 |
|  | UniBa | 12 | 0.78 | 0.91 | 0.82 | 0.37 |
|  | UniBa | 28 | 0.85 | 0.89 | 0.94 | 0.90 |
|  | UniBa | 52 | 0.75 | 0.95 | 0.98 | 0.30 |
| **Diaphragm twitch HRT (ms)** | UniBa | 8 | 0.94 | 0.96 | 0.48 | 0.16 |
|  | UniBa | 12 | 0.14 | 0.99 | 0.59 | 0.99 |
|  | UniBa | 28 | 0.18 | 0.63 | 0.59 | 0.26 |
|  | UniBa | 52 | **0.025** | 0.31 | 0.81 | 0.92 |
| **Diaphragm twitch specific force (kN/m^2^)** | UniBa | 8 | 0.26 | 0.06 | **0.001** | 0.06 |
|  | UniBa | 12 | **0.013** | **0.004** | 0.09 | **0.007** |
|  | UniBa | 28 | **0.001** | 0.13 | 0.61 | 0.17 |
|  | UniBa | 52 | **<0.001** | **0.026** | 0.11 | 0.35 |
| **Diaphragm tetanic specific force (kN/m^2^)** | UniBa | 8 | 0.32 | 0.17 | 0.49 | 0.25 |
|  | UniBa | 12 | **0.049** | **<0.001** | 0.06 | 0.20 |
|  | UniBa | 28 | **<0.001** | **0.002** | 0.17 | 0.30 |
|  | UniBa | 52 | **<0.001** | **0.010** | 0.14 | 0.99 |
| **Diaphragm force drop at 5^th^ pulse (%)** | UniBa | 8 | 0.87 | 0.11 | 0.28 | 0.97 |
|  | UniBa | 12 | **0.030** | 0.25 | 0.37 | 0.97 |
|  | UniBa | 28 | **<0.001** | **<0.001** | 0.39 | 0.47 |
|  | UniBa | 52 | 0.99 | 0.15 | 0.35 | 0.99 |
| **Diaphragm force drop at 10^th^ pulse (%)** | UniBa | 8 | 0.97 | 0.11 | 0.30 | 0.93 |
|  | UniBa | 12 | **0.003** | 0.05 | 0.60 | 0.83 |
|  | UniBa | 28 | **<0.001** | **<0.001** | 0.98 | 0.71 |
|  | UniBa | 52 | 0.95 | **0.016** | 0.24 | 0.99 |
| **Diaphragm force recovery at 5 min (%)** | UniBa | 8 | 0.67 | 0.31 | 0.76 | 0.44 |
|  | UniBa | 12 | 0.10 | **0.016** | 0.99 | 0.99 |
|  | UniBa | 28 | **<0.001** | **0.013** | 0.90 | 0.99 |
|  | UniBa | 52 | 0.09 | **0.002** | 0.91 | 0.99 |
| **Diaphragm force recovery at 15 min (%)** | UniBa | 8 | 0.68 | 0.96 | 0.15 | 0.56 |
|  | UniBa | 12 | 0.24 | **0.005** | 0.98 | 0.99 |
|  | UniBa | 28 | **0.003** | **0.015** | 0.99 | 0.02 |
|  | UniBa | 52 | **0.022** | **<0.001** | 0.95 | 0.99 |

**Supplementary Table 13. Strain comparisons of *ex vivo* physiology of the diaphragm.** Adjusted *P­*-values of the strain by strain comparisons for each of the study sites. *P*<0.05 is indicated in bold.

| **Outcome** | **Site** | **Age**  **(weeks)** | **BL10-WT vs.**  **BL10-*mdx*** | **D2-WT**  **vs. D2-*mdx*** | **BL10-*mdx***  **vs. D2-*mdx*** | **BL10-WT**  **vs. D2-WT** |
| --- | --- | --- | --- | --- | --- | --- |
| CK (U/L) | LUMC | 2 | **0.020** | 0.14 | 0.86 | 0.80 |
|  | UniBa | 4 | 0.46 | 0.41 | 0.99 | 0.99 |
|  | LUMC | 4 | **0.003** | 0.07 | 0.41 | 0.99 |
|  | UniBa | 8 | **<0.001** | 0.33 | 0.13 | 0.46 |
|  | LUMC | 8 | **0.002** | **0.027** | 0.91 | 0.98 |
|  | UniBa | 12 | **0.004** | **0.001** | 0.98 | 0.07 |
|  | LUMC | 12 | **0.019** | **0.004** | 0.09 | 0.55 |
|  | UniBa | 28 | **<0.001** | **0.030** | **<0.001** | 0.99 |
|  | LUMC | 28 | **0.040** | 0.09 | **0.014** | 0.99 |
|  | UniBa | 52 | **<0.001** | 0.67 | **0.007** | 0.90 |
|  | LUMC | 52 | 0.33 | 0.89 | 0.39 | 0.83 |
| LDH (U/L) | UniBa | 4 | 0.94 | 0.30 | 0.59 | 0.99 |
|  | UniBa | 8 | **0.011** | 0.61 | 0.06 | 0.94 |
|  | UniBa | 12 | 0.90 | 0.30 | 0.99 | 0.54 |
|  | UniBa | 28 | **0.004** | **0.010** | 0.22 | 0.74 |
|  | UniBa | 52 | **0.016** | 0.99 | 0.10 | 0.81 |

**Supplementary Table 14. Strain comparisons of plasma CK and LDH levels.** Adjusted *P­*-values of the strain by strain comparisons for each of the study sites. *P*<0.05 is indicated in bold. CK; creatine kinase, LDH; lactate dehydrogenase.
